# Neuronal Cell-Cycle Re-entry Defines Divergent Outcomes Through Replication-Dependent DNA Damage in ALS

**DOI:** 10.64898/2026.02.13.705790

**Authors:** Jonathan Plessis-Belair, Roger B. Sher

**Author notes:** **Corresponding authors:** Roger Sher.

## Abstract

Cell-cycle dysregulation has emerged as a shared mechanism of neuronal loss across neurodegenerative diseases (NDDs), including amyotrophic lateral sclerosis (ALS), Alzheimer’s disease, and Parkinson’s disease. In post-mitotic neurons, aberrant reactivation of cell-cycle signaling precedes degeneration, yet the upstream triggers and functional consequences of this process remain poorly defined. Nucleocytoplasmic transport (NCT) dysfunction, a hallmark of ALS and related disorders, disrupts the spatial distribution of key regulatory proteins and may contribute to maladaptive cell-cycle activation. Our recent evidence suggests that impaired nuclear import may initiate, rather than merely accompany, neuronal cell-cycle re-entry. Here, we show that cell-cycle activation in motor neurons distinguishes molecular subtypes and outcomes in ALS. We analyzed the AnswerALS transcriptomic cohort and identified a patient cluster characterized by robust upregulation of cyclins B and D. Clusters with lower levels of cell-cycle gene expression exhibited accelerated ALSFRS-R decline, whereas the highest cyclin-expressing cluster demonstrated comparatively improved functional trajectories over time. To test whether NCT disruption can mechanistically drive aberrant cell-cycle activation, we pharmacologically inhibited importin-β in human iPSC-derived spinal motor neurons. NCT disruption induced widespread proteomic mislocalization, including TDP-43 pathology, and triggered a transient wave of cell-cycle activity preceding neuronal death. Mechanistically, we identified DNA-replication initiation as a pathological event driving degeneration and demonstrated that selective inhibition of G1/S-associated CDK4/6 activity confers neuroprotection. Together, these findings link impaired nuclear import to maladaptive cell-cycle reactivation in neurons and highlight stage-specific engagement of the cell-cycle machinery as a determinant of neuronal vulnerability in ALS.

## Background

Cell cycle dysregulation has emerged as a unifying feature of neuronal loss across multiple neurodegenerative diseases (NDDs) (*1*), including Amyotrophic Lateral Sclerosis (ALS)(*2–4*), Alzheimer’s disease (AD)(*5–9*), and Parkinson’s disease (PD)(*8, 10–13*). These events in post-mitotic neurons are characterized by the re-expression of signaling effectors traditionally associated with cell cycle progression and proliferation (*14, 15*). Although only a small subset of neurons exhibit neuronal cell cycle re-entry at any given time, the frequency of these events increases from early to late disease stages, suggesting a threshold-dependent activation mechanism that may precede overt neurodegeneration (*14*).

The upstream mechanisms that trigger neuronal cell cycle activation remain poorly understood. In ALS, nucleocytoplasmic transport (NCT) dysfunction is a major pathogenic process, reflecting a progressive decline in the efficiency of protein transport between nuclear and cytoplasmic compartments (*16–19*). Disruption of this transport balance leads to widespread protein mislocalization and aggregation, implicating both nuclear import and export pathways in the pathogenesis of NDDs (*20–24*). Notably, TAR DNA-binding protein 43 (TDP-43), a defining pathological hallmark of ALS, is directly mislocalized in the context of NCT impairment (*16*). Furthermore, we and others have shown that these NCT defects are an upstream driver of cell cycle activation (*25, 26*), supporting a model in which impaired transport dynamics may actively contribute to, rather than simply reflect, neurodegenerative progression.

Despite these observations, the mechanisms by which NCT disruption and aberrant cell cycle activation ultimately lead to neuronal death remains unclear. Notably, cell-cycle reactivation in post-mitotic neurons has been proposed as a stress-responsive attempt to engage DNA damage repair pathways (*14, 27*). In this context, limited activation of cell-cycle machinery may represent an adaptive response to genomic instability. However, failure to resolve this response, particularly progression toward DNA replication, has frequently been associated with neuronal dysfunction and cell death (*28*). Complicating this picture is the multifunctionality of cell cycle regulators in neurons. Evolutionary constraints have repurposed these molecules for roles in dendritic and axonal maintenance, where they support normal physiology but may become deleterious under stress or mislocalization (*1, 29, 30*). Thus, understanding neuronal vulnerability requires moving beyond traditional mitotic frameworks to integrate the spatial, temporal, and contextual dynamics of cell-cycle proteins within the post-mitotic neuronal environment.

Here, we investigated cell-cycle transcriptomic signals in the AnswerALS cohort and how pharmacological impairment of nuclear import reshapes neuronal cell-cycle dynamics and survival in vitro. Through transcriptomic analysis of the AnswerALS cohort, we identified a molecular subgroup characterized by robust upregulation of cyclins B and D. Clusters with lower levels of cell-cycle gene expression were associated with accelerated ALSFRS-R decline, whereas the highest cyclin-expressing cluster demonstrated comparatively improved functional trajectories, suggesting that the magnitude of cyclin activation may reflect distinct neuronal trajectories. To test whether nucleocytoplasmic transport (NCT) disruption mechanistically drives these changes *in vitro*, we pharmacologically inhibited importin-β-mediated nuclear import in human iPSC-derived spinal motor neurons, which induced widespread protein mislocalization characteristic of ALS pathology (*31*). We found that aberrant cell-cycle activation emerges as a downstream response to NCT dysfunction and precedes neuronal death. Mechanistically, our data implicate S-phase entry, marked by neuronal DNA replication initiation, as a critical pathological transition, and demonstrate that restricting G1/S-associated CDK4/6 activity confers neuroprotection. Together, these findings redefine neuronal cell-cycle reactivation as a stage-specific process in which progression toward replication, rather than initial activation alone, determines vulnerability. More broadly, this work highlights the importance of capturing early molecular events, as late-stage post-mortem analyses may preferentially reflect surviving neuronal populations rather than those that have undergone degeneration.

## Results

### Cell cycle are-entry defines a distinct subgroup of ALS patients

Given the extensive molecular heterogeneity reported in ALS, we first performed an unbiased transcriptomic stratification of iPSC-patient-derived motor neurons from the AnswerALS cohort (25). From the full registry, we selected 624 individuals (445 ALS patients, 161 healthy controls, 10 non-ALS motor neuron disease [MND] cases, and 8 asymptomatic ALS gene carriers) for whom high-quality RNA-sequencing data were available (Fig. 1A, Table S1). Unsupervised clustering using UMAP dimensionality reduction followed by k-means clustering (k = 10) identified several distinct molecular subgroups across the cohort (Fig. 1B, Table S2). Among these, one subgroup (cluster 7) was defined by robust upregulation of cell cycle–associated genes, including cyclins D (*CCND1*), A (*CCNA2*), and B (*CCNB1*), but notably not cyclin E (*CCNE1*) (Fig. 1C-E). Ranking differentially expressed genes by Wald statistic revealed that DNA maintenance programs and cell cycle pathways are among most enriched across both Reactome and KEGG annotations (Fig. S1A-B). Directional enrichment analysis further showed that these cell cycle pathways were concentrated within the upregulated gene set (Fig. S1C-D), whereas neuronal, synaptic, and neurodegenerative pathways were preferentially enriched among downregulated genes (Fig. S1E-F). Together, these findings indicate that transcriptional signatures of cell-cycle activation define a distinct subset of ALS patients within the AnswerALS cohort.

**Figure 1:**
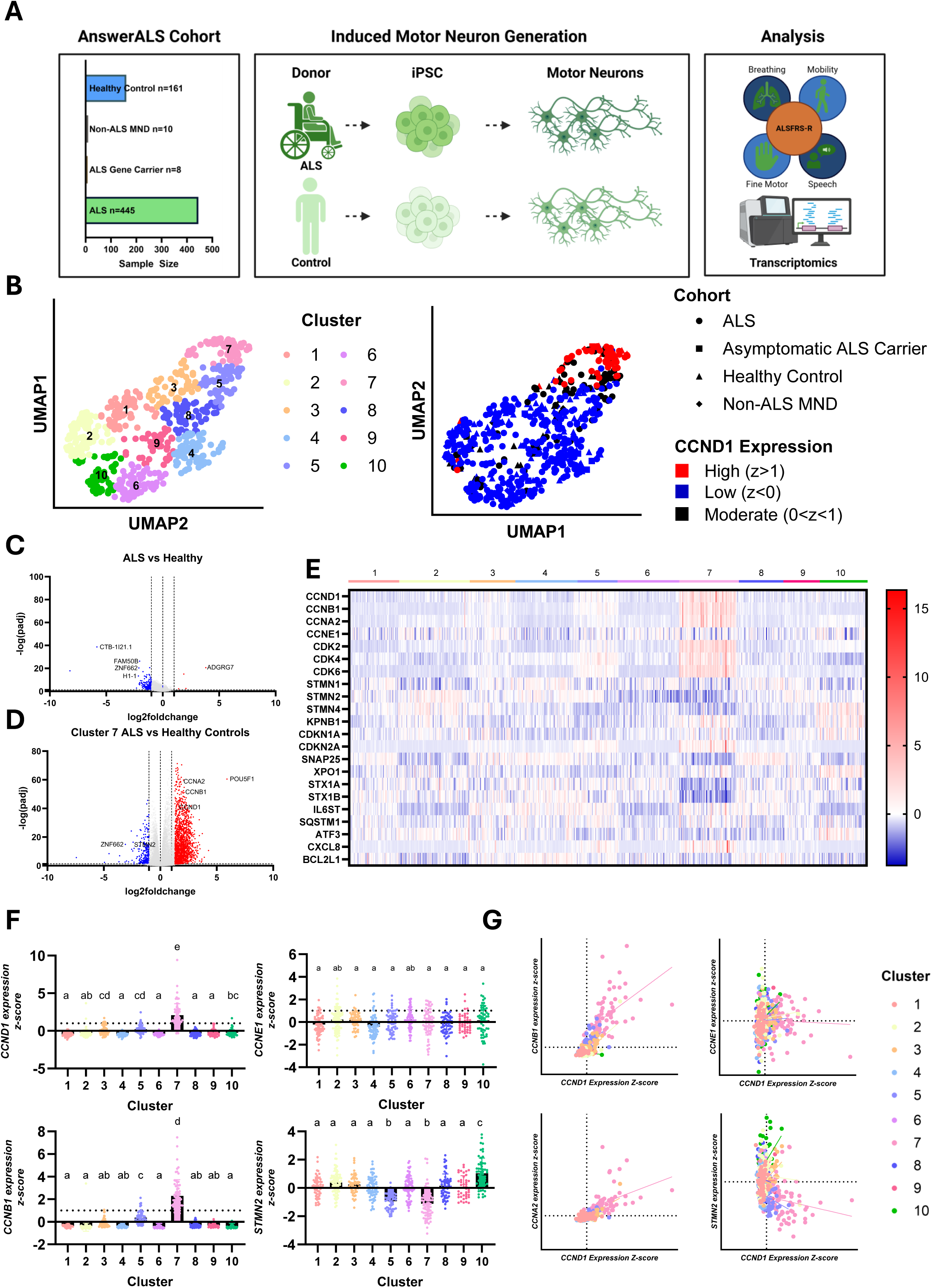
Cell cycle activation defines a distinct subgroup of ALS patients. **(A)** Schematic of the AnswerALS cohort (161 healthy controls, 10 non-ALS motor neuron disease, 8 asymptomatic ALS gene carriers, and 445 ALS patients) and pipeline of patient iPSC-derived motor neurons with clinical ALSFRS-R progression scores and transcriptomic analyses. **(B)** UMAP analysis of 624 patient RNA-seq samples with k-means clustering (k=10), highlighting cluster 7 as enriched for high *CCND1* expression (z > 1). **(C–D)** Volcano plots comparing ALS patients (n=445) versus healthy controls (n=161) (C), and cluster 7 ALS patients (n=69) versus healthy controls (n=161) (D). **(E)** Heatmap of selected cell-cycle and disease-relevant gene expression z-scores organized by patient cluster. **(F)** Pairwise comparisons of selected genes (*CCND1, CCNB1, CCNE1,* and *STMN2*) across patient clusters. Significance is indicated by letter groupings and was determined by one-way ANOVA with Tukey’s multiple-comparison test (p < 0.05). **(G)** Correlation plots of *CCND1* z-scores with selected genes (*CCNB1, CCNA2, CCNE1,* and *STMN2*) across patient clusters with simple linear regressions. Bar graphs represent mean ± standard deviation. All data points and statistical comparisons are provided in the Source Data file. Schematics created with BioRender.com.

To further characterize transcriptional differences across clusters, we performed pairwise comparisons of representative genes associated with cell cycle activation (*CCND1*, *CCNB1*, *CCNE1*) and neuronal integrity (*STMN2*) (Fig 1F). *CCND1* and *CCNB1* expression were markedly enriched in cluster 7, with minimal expression in other clusters, whereas *STMN2* expression was reduced in clusters 5 and 7 (Fig 1F). In contrast, *CCNE1* expression did not vary significantly across clusters (Fig 1F). To assess relationships between these gene programs, we next examined correlations between *CCND1* and other markers (Fig 1G). *CCND1* expression positively correlated with *CCNB1* and *CCNA2*, consistent with coordinated activation of G1 and G2 cell cycle regulators, while showing no correlation with *CCNE1* (S phase) and a strong negative correlation with *STMN2* (Fig 1G).

The enrichment of cell cycle-associated transcripts in this subgroup suggests that aberrant reactivation of cell cycle pathways may occur in a subset of ALS patients, or at a particular stage of disease progression. Such activation has been previously linked to neuronal stress responses and vulnerability (*27, 32*), raising the possibility that these patients represent a molecularly distinct disease subtype. Notably, the lack of S-phase associated cyclin E upregulation implies partial or selective engagement of the cell cycle machinery rather than full cell cycle re-entry, consistent with non-proliferative cell cycle activation observed in neuronal models (*33*).

### Differential expression of cell cycle effectors in ALS patients defines disease progression

Having identified molecularly distinct subgroups of ALS patients, including one defined by cell cycle activation, we next examined whether these clusters correspond to differences in disease outcome. We first assessed overall survival across all ten transcriptomic clusters using Kaplan-Meier analysis. This analysis revealed no statistically significant difference in survival probability between clusters (Fig. 2A), indicating that the observed transcriptional heterogeneity is not associated with survival within the current follow-up period. Subsequently, there was so significant differences between the age of disease onset or the age at time of death between patients across clusters (Table S2). However, given that many individuals remain alive, and survival follow-up is still ongoing, these findings do not exclude potential long-term survival differences.

**Figure 2:**
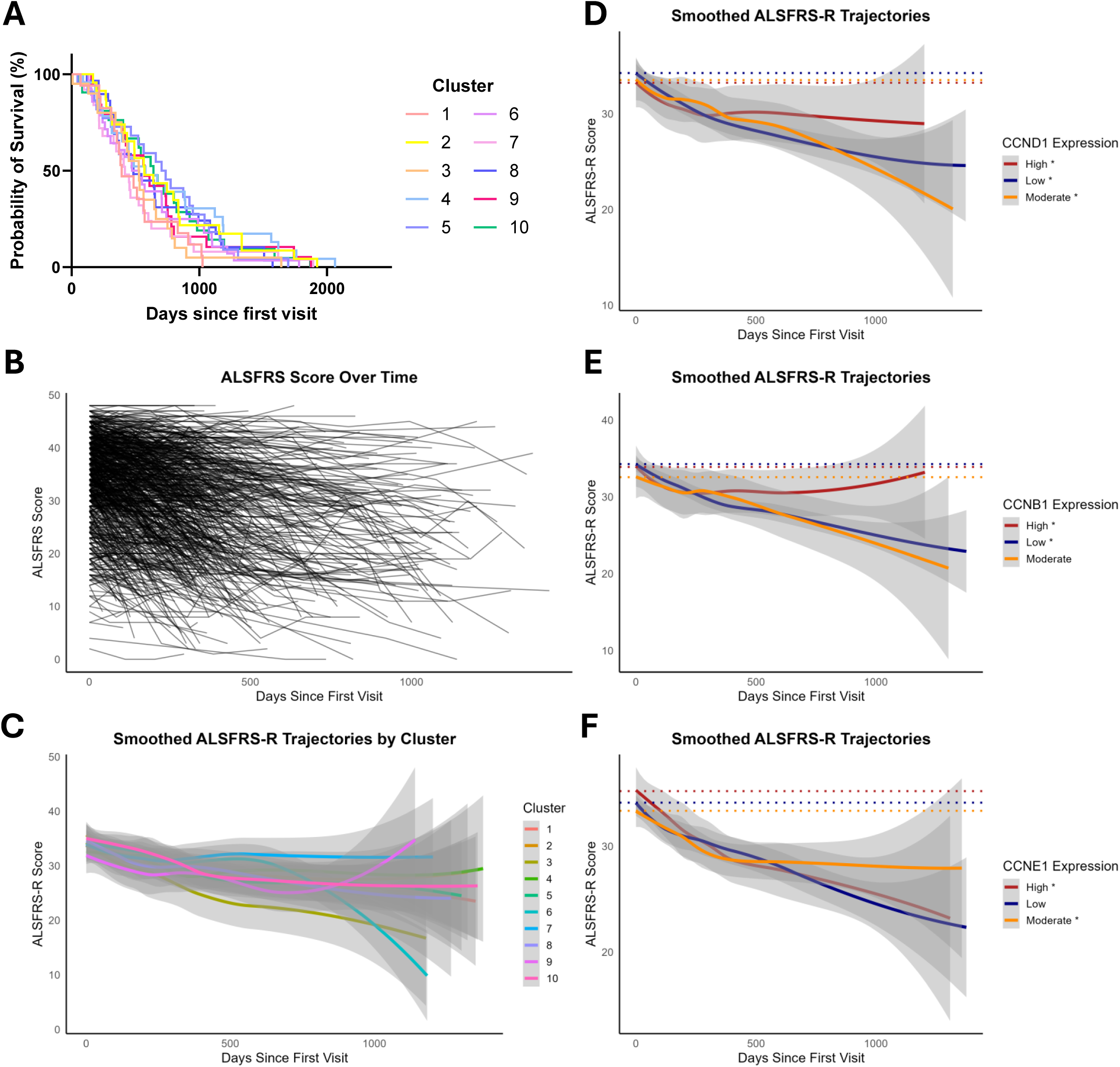
Levels of Cell Cycle Activation in ALS patients defines disease progression. **(A)** Kaplan–Meier survival analyses across ALS patient clusters (n=227), with significance assessed using the log-rank (Mantel–Cox) test. **(B)** ALSFRS-R score trajectories from the first recorded patient visit (n=290). **(C)** Smoothed ALSFRS-R trajectories across ALS patient clusters (n=290), with group differences evaluated using linear mixed-effects models. Pairwise cluster comparisons: 1-8**, 2-7*, 2-8*, 3-7*, 4-8**, 5-6**, 5-8**, 6-8**, 7-8***, 8-9**, 8-10**. **(D–F)** Smoothed ALSFRS-R trajectories for *CCND1* (D), *CCNB1* (E), and *CCNE1* (F) across high (z > 1), moderate (0 < z < 1), and low (z < 0) gene-expression groups (n = 290), analyzed using linear mixed-effects models. **(G)** All statistical comparisons are provided in the Source Data file. Significance values are indicated as * p<0.05, ** p<0.01, *** p<0.001.

We next evaluated disease severity and functional decline using the ALS Functional Rating Scale-Revised (ALSFRS-R). Examination of clinical metrics across transcriptional subgroups revealed distinct functional trajectories among ALS patient clusters. Cluster 7, defined by high expression of cell cycle effectors (Fig 1), exhibited the slowest disease progression (Fig 2C). In contrast, clusters 6 and 9, which display only moderate expression of cell cycle effectors, showed divergent outcomes: cluster 6 declined steadily, whereas cluster 9 demonstrated a positive increase later during progression. To further explore this relationship, patients were stratified by the degree of cell cycle activation. Individuals with high expression of cell cycle-associated genes maintained the highest ALSFRS-R scores throughout the disease course, while those with moderate activation demonstrated the lowest scores, suggesting a biphasic relationship between cell cycle activation and functional decline (Fig 2D-F).

Interestingly, the relationship between cell-cycle activation and disease severity was both gene-specific and phase-dependent. Elevated expression of *CCND1* and *CCNB1*, corresponding to G1 and G2 phase regulators, respectively, was associated with improved functional outcomes (Fig 2D, E). In contrast, both high and low expression of *CCNE1* (S-phase) correlated with reduced ALSFRS-R scores, whereas moderate *CCNE1* expression was linked to the most favorable prognosis (Fig 2F). Among the cyclins analyzed, CCNB1 showed the strongest association with improved survival, with mean ALSFRS-R scores returning toward baseline over time. Collectively, these findings indicate that phase-specific reactivation of the cell cycle exerts distinct effects on disease trajectory, suggesting that controlled or partial engagement of specific cell-cycle modules may differentially influence neuronal function and survival.

To account for potential confounding variables that might influence disease severity independently of transcriptional state, we evaluated sex, site of onset, and clinical mutation status across ALS patient clusters (Fig S2A-D). Neither sex nor clinical mutation category showed statistically significant differences in ALSFRS-R progression rates, although individual mutations displayed substantial variability in disease course (Fig S2E-F). In contrast, site of onset exerted the strongest effect on functional decline, with markedly different ALSFRS-R slopes across onset categories (Fig S2G). This effect was most pronounced in cluster 7: patients with bulbar or multiple sites of onset exhibited accelerated decline, whereas those with limb onset or no documented site of onset showed minimal change over time (Fig S2G).

We next examined past medical history as an additional confounder. Medical conditions were grouped into major categories (cardiovascular, metabolic, surgical, gastrointestinal, urologic, neurologic, etc.) based on available diagnoses, with rare or unclassifiable entries grouped into an “other” category (Fig S2H, Source Data). Cluster-level stratification revealed increased prevalence of metabolic, gastrointestinal, urologic, and neurologic comorbidities within cluster 7 (Fig S2I). Notably, patients with a metabolic disease history (e.g., obesity, dyslipidemia, diabetes) showed significantly slower ALSFRS-R decline, consistent with previously reported protective effects of metabolic dysfunction in ALS (Fig S2J) (*34*). However, this benefit was attenuated within cluster 7, suggesting a more complex interaction between metabolic disease, cell-cycle activation, and neuroprotection. In contrast, neurological comorbidities (e.g., brain trauma, mild cognitive impairment, spinal injury) were associated with accelerated disease progression, an effect that remained within cluster 7 (Fig S2K). Given the established connection between cell-cycle dysregulation and cancer biology, we also examined whether a history of cancer was linked to disease course in cluster 7; however, no differences in ALSFRS-R progression were detected aside from a reduction in overall lifespan (Fig S2L).

Together, these analyses reveal that clinical and medical comorbidities exert heterogeneous effects on ALS disease course, with site of onset and metabolic or neurologic history emerging as the strongest modifiers of progression. Importantly, several of these effects are amplified or altered within cluster 7, the subgroup defined by high cell-cycle activation, suggesting that transcriptional state interacts with clinical context to shape disease trajectory. These findings highlight the importance of integrating molecular stratification with patient-specific clinical variables when interpreting ALS heterogeneity and identifying factors that modulate progression.

### Importin-β inhibition disrupts nuclear import and induces ALS-associated TDP-43 mislocalization in human iPSC-derived motor neurons

Building on our patient-based findings, we next investigated the mechanistic relationship between nuclear import defects and cell cycle dysregulation using a human iPSC-derived motor neuron model. We employed a doxycycline-inducible *NGN2* overexpression iPSC line (*35–37*), which markedly accelerates neuronal maturation and enables the generation of spinal motor neurons within a short timeframe (Fig. 3A). To account for variability across differentiations, each iPSC line was independently differentiated at least three times. The resulting cultures consistently yielded ∼73–90% ChAT⁺/MAP2⁺ neurons with minimal S100β⁺ contamination (<3%) (Fig. 3B-C). Expression analysis further confirmed robust induction of spinal motor neuron markers, including *HOXB4/C6*, *MNX1*, and *ChAT* (Fig. 3D).

**Figure 3:**
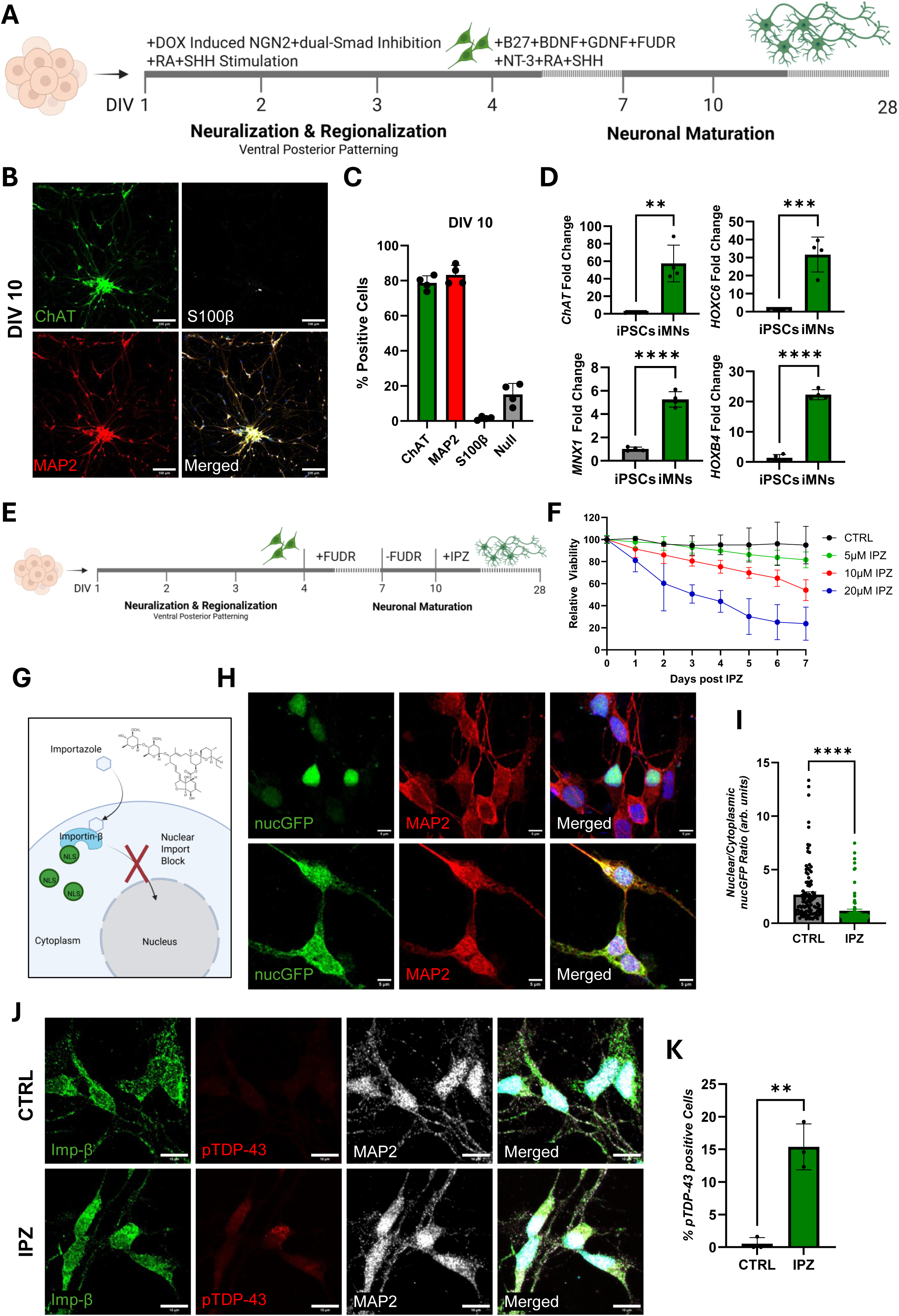
Importin-β inhibition disrupts nuclear import and induces ALS-associated TDP-43 mislocalization in human iPSC-derived motor neurons. **(A)** Timeline of doxycycline-inducible NGN2 iPSC-derived motor neuron generation and maturation. **(B)** Representative immunostains of ChAT (green), MAP2 (red), and S100β (white) in iPSC-derived motor neurons at DIV10. DAPI (blue) is a nuclear counterstain in the merged image. **(C)** Quantification of the percentage of ChAT+, MAP2+, S100β+, or null (DAPI-only) cells from (B) (n=4 replicates). **(D)** qPCR quantification of spinal motor neuron markers *ChAT, MNX1,* and *HOXC6/B4,* shown as fold change relative to undifferentiated iPSCs, analyzed by unpaired two-tailed t-tests (n=4 replicates). **(E)** Timeline of iPSC-derived motor neuron generation treated with the importin-β antagonist importazole (IPZ) at DIV10. **(F)** Seven-day viability curves of DMSO control and IPZ-treated (5 μM, 10 μM, and 20 μM) motor neurons, analyzed by two-way ANOVA with Tukey’s multiple-comparison test (p < 0.05) (n=3 replicates). **(G)** Schematic of importazole inhibition of importin-β-mediated nuclear import. **(H)** Representative immunostains of GFP-3X-NLS (green) and MAP2 (red) in transfected iPSC-derived motor neurons at DIV10. DAPI (blue) is a nuclear counterstain in the merged image. **(I)** Quantification of nuclear/cytoplasmic intensity ratios nucGFP (H), analyzed using an unpaired two-tailed t-test (p < 0.05) (n=3 replicates). **(J)** Representative immunostains of importin-β (green), pTDP-43 (red), and MAP2 (white) in iPSC-derived motor neurons at DIV10. DAPI (blue) is a nuclear counterstain in the merged image. **(K)** Quantification of the percentage of pTDP-43–positive motor neurons from (J), analyzed using an unpaired two-tailed t-test (p < 0.05) (n=3 replicates). Bar graphs represent mean ± standard deviation and significance values are indicated as * p<0.05, ** p<0.01, *** p<0.001. Scale bars: 100 μm (B) and 10 μm (G–I). Replicates are independent differentiations to account for batch variability. Schematics created with BioRender.com. All data points and statistical comparisons are provided in the Source Data file.

Using this model, we implemented a drug-inducible paradigm in which importazole, a selective inhibitor of importin-β–mediated nuclear import, was used to recapitulate key aspects of ALS-associated nuclear transport defects (Fig 3E) (*25*). This system provided precise temporal control of pathological processes, enabling direct investigation of how impaired nuclear import influences neuronal homeostasis and cell cycle reactivation. To determine the optimal conditions for modeling ALS-associated pathology, we assessed the effect of importazole concentration on motor neuron viability. Dose-response curves to a 2 day importazole treatment of iPSC-derived motor neurons showed sublethal concentrations below 10 µM (Fig S4). Using these values, we assessed the time-dependent cell survival of iPSC-derived motor neurons with varying concentrations of importazole (5, 10, and 20 µM) (Fig 3F). Treatment with 20 µM importazole resulted in rapid and extensive neuronal loss, with a large majority of cells (∼50%) dying within 2–3 days (Fig 3F). In contrast, 5 µM treatment had minimal impact on neuronal viability, showing little to no cell death across the experimental timeframe (Fig 3F). The intermediate concentration of 10 µM induced a modest and progressive decline in viability, without triggering an overt cell death response (Fig 3F). These observations establish 10 µM as a sublethal, disease-relevant dose suitable for subsequent mechanistic investigations.

We next confirmed that importazole effectively blocks nuclear import in human spinal motor neurons using a nuclear GFP reporter (3× SV40 NLS) expressed under the human synapsin (hSyn) promoter (*38*) (Fig. 3G). Importazole robustly impaired nuclear accumulation of nucGFP across the majority of neurons, validating the population-wide inducibility of this model (Fig. 3H-I). (*38*)We then asked whether nuclear import disruption also elicits hallmark features of ALS pathology. Immunostaining for phosphorylated TDP-43 (pTDP-43), a pathological species observed in approximately 97% of ALS cases (*39*), revealed significant cytoplasmic aggregation in 30-40% of treated motor neurons (Fig. 3J-K). These findings demonstrate that pharmacological inhibition of importin-β not only recapitulates nuclear import defects but also drives ALS-like cytoplasmic TDP-43 pathology in human spinal motor neurons.

### SILAC-based spatial proteomics reveals widespread nuclear-cytoplasmic redistribution following importin-β inhibition

To comprehensively define the molecular consequences of importazole-induced nuclear import disruption, we employed Stable Isotope Labeling in Cell Culture (SILAC) combined with nuclear-cytoplasmic fractionation to map subcellular proteomic redistribution (Fig. 4A) (34, 35). This approach enables quantitative assessment of protein localization shifts between compartments under basal and pathological conditions.

**Figure 4:**
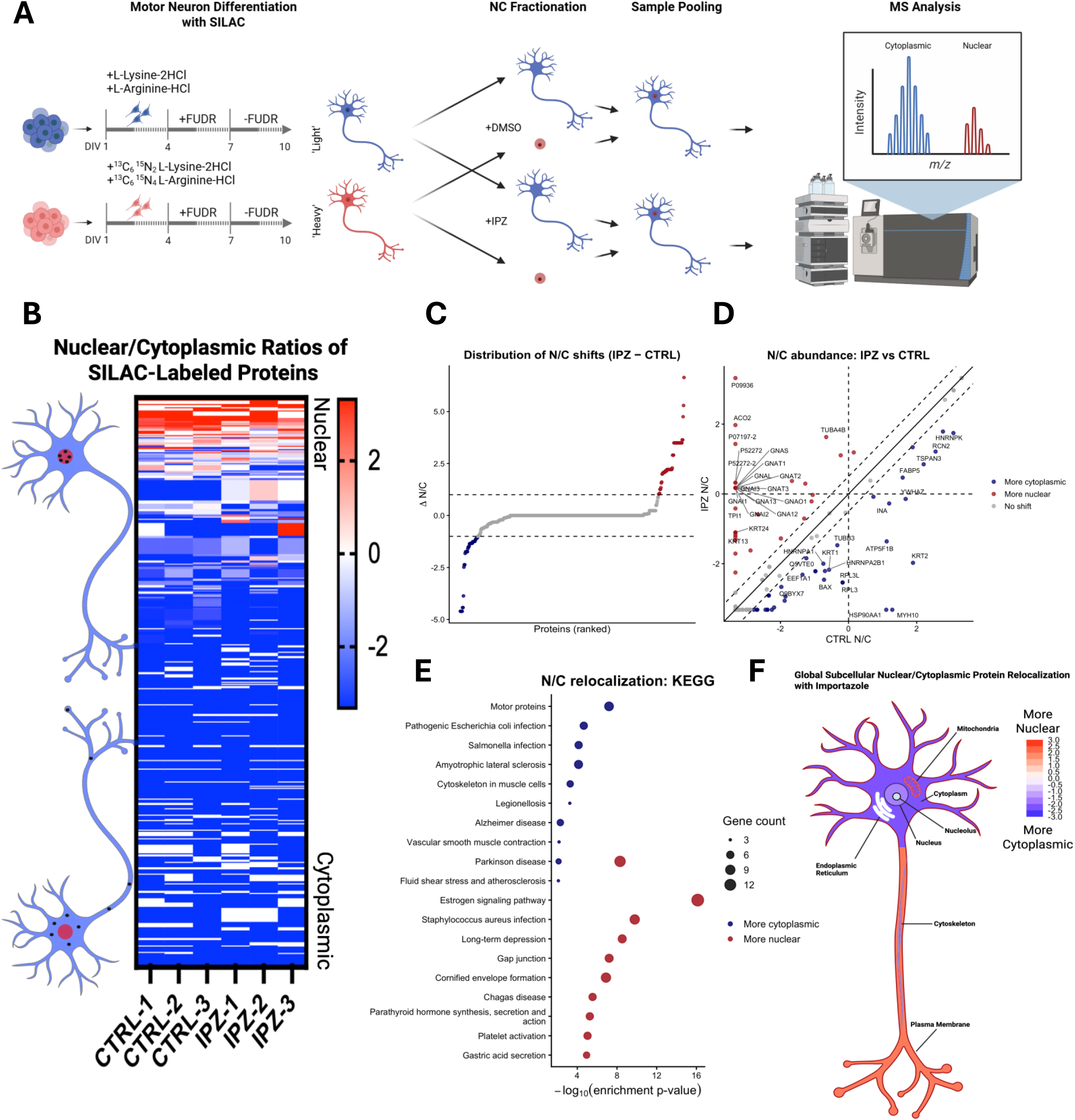
SILAC-based spatial proteomics reveals widespread nuclear–cytoplasmic redistribution following importin-β inhibition. **(A)** Schematic illustrating incorporation of heavy and light labeled amino acids in iPSC-derived motor neurons, followed by importazole treatment, subcellular fractionation, and LC-MS/MS analysis of nuclear (heavy) and cytoplasmic (light) fractions. **(B)** Heatmap depicting log₂ fold-change nuclear-to-cytoplasmic (N/C) ratios of detected peptides, with nuclear enrichment shown in red, cytoplasmic enrichment in blue, and evenly distributed proteins in white. (n=3 replicates; Source Data) **(C)** Waterfall plot of differential N/C ratios (IPZ–CTRL), highlighting proteins with increased nuclear localization (red; >0.5) or increased cytoplasmic localization (blue; <−0.5) following importazole treatment. **(D)** Scatter plot showing the distribution of log₂ fold-change N/C ratios under CTRL and importazole (IPZ) conditions. **(E)** KEGG pathway enrichment analysis of nuclear- and cytoplasmic-enriched differentially localized proteins following importazole treatment. **(F)** Schematic summarizing global protein relocalization in iPSC-derived motor neurons after importazole treatment, with proteins shifting toward the nucleus shown in red and toward the cytoplasm shown in blue. Replicates are independent differentiations to account for batch variability. Schematics created with BioRender.com. All data points and statistical comparisons are provided in the Source Data file.

Because SILAC requires near-complete incorporation of labeled amino acids, cells must undergo at least three passages. We therefore first optimized this strategy using the mitotic neuronal cell line SK-N-MC, which allowed for robust labeling efficiency across the proteome following importazole-mediated inhibition of importin-β (Fig. S5). Global changes in protein localization were visualized by a heatmap of log₂ nuclear-to-cytoplasmic fold changes, with blue indicating cytoplasmic enrichment, red indicating nuclear enrichment, and white denoting equal distribution (Fig. S5A). Fractionation quality was confirmed by strong nuclear enrichment of histones and lamins and cytoplasmic enrichment of actins and tubulins (Fig. S5B), validating the robustness of the spatial proteomic workflow.

Using this validated system, we next identified proteins exhibiting significant subcellular redistribution following importazole treatment (Fig. S5C). Reactome pathway analysis revealed that cytoplasmically mislocalized proteins were strongly enriched for cell cycle-related pathways, whereas nuclear-enriched proteins were associated with translation, axon guidance, and nervous system development (Fig. S5D). Complementary KEGG analysis showed significant enrichment of neurodegenerative disease pathways in both fractions, including ALS, Parkinson’s disease, prion disease, and Alzheimer’s disease, with ALS emerging as the most highly enriched pathway.

We then extended this approach to iPSC-derived motor neurons, which required incorporation of heavy and light isotopes at the iPSC stage and maintenance throughout neuronal differentiation (Fig. 4A). Although fewer labeled peptides were detected following neuronal maturation, the relative distribution of nuclear and cytoplasmic proteins remained consistent (Fig. 4B, S5E). Importazole treatment induced bidirectional protein redistribution, with comparable numbers of proteins mislocalized to the cytoplasm and nucleus, indicating that impaired nuclear import disrupts global nucleocytoplasmic organization rather than causing simple cytoplasmic accumulation (Fig. 4C-D, S5F). Notably, cytoplasmically mislocalized proteins included HNRNPA1, HNRNPA2B1, HSP90AA1, and BAX, factors previously implicated in ALS and related neurodegenerative diseases (*40–43*) (Fig. 4D).

Consistent with these observations, KEGG pathway analysis of redistributed proteins in motor neurons highlighted ALS, Alzheimer’s disease, and Parkinson’s disease as highly enriched among cytoplasmically shifted proteins (Fig. 4E). In contrast to the mitotic cell line, Reactome analysis in motor neurons revealed enrichment of neuronal pathways, including cell junction organization and neurotransmitter receptor signaling (Fig. S5G), underscoring cell-type-specific responses to nuclear import disruption. Further stratification by predicted functional localization showed that mitochondrial, cytoskeletal, and plasma membrane-associated proteins preferentially redistributed to the nucleus, whereas nuclear, nucleolar, and cytoplasmic proteins accumulated in the cytoplasm following importazole treatment (Fig. 4F).

Collectively, these data demonstrate that pharmacological inhibition of importin-β disrupts nucleocytoplasmic organization in both mitotic neuronal cells and post-mitotic iPSC-derived motor neurons, driving widespread protein mislocalization that converges on neurodegenerative disease pathways, most prominently ALS.

### Nuclear import inhibition triggers cell-cycle stage-specific re-entry in motor neurons

Given the strong enrichment of cell cycle-related pathways identified in our spatial proteomics analysis, we next examined whether importazole-induced nuclear import disruption triggers cell cycle activation in human spinal motor neurons. Immunostaining for the proliferation marker Ki-67 was performed across a six-day time course following importazole treatment (Fig. 5A-C). We observed a marked increase in Ki-67–positive neurons as early as day 1, with elevated levels persisting through day 3. By days 4-6, Ki-67 expression was variable and rare, suggesting that neurons exhibiting early cell cycle re-entry may subsequently undergo cell death.

**Figure 5:**
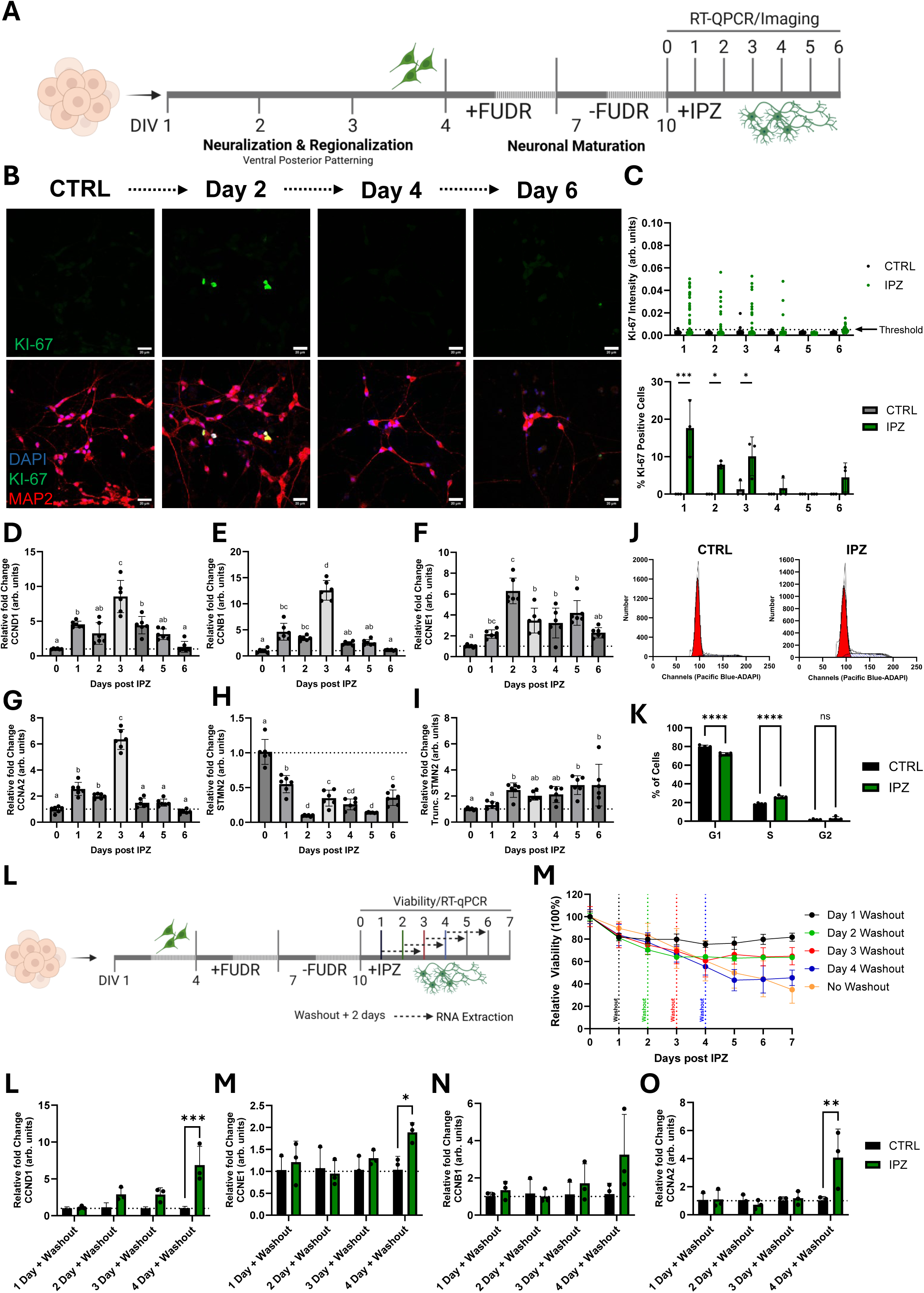
Nuclear import inhibition triggers cell-cycle stage-specific re-entry in motor neurons. **(A)** Timeline of iPSC-derived motor neuron generation treated with the importin-β antagonist IPZ at DIV10 and time-course of associated RT-qPCR and imaging analysis. **(B)** Representative immunostains of KI-67 (green) in iPSC-derived motor neurons at indicated timepoints post IPZ treatment (CTRL [Day 1] and IPZ Day 2, 4, and 6). DAPI (blue) and MAP2 (red) is a nuclear and neuronal counter stain in the merged image. **(C)** Quantification of KI-67 intensity and percentage of KI-67 (mean intensity >0.01 threshold) positive motor neurons, analyzed by two-way ANOVA with Tukey’s multiple-comparison test (p < 0.05) (n=3 replicates; Source Data). **(D-I)** qPCR quantification of cell-cycle and disease-relevant genes (*CCND1, CCNB1, CCNE1, CCNA2, STMN2*, and truncated *STMN2*) at days 0-6 post IPZ treatment, shown as fold change relative to control. Significance is indicated by letter groupings and was determined by one-way ANOVA with Tukey’s multiple-comparison test (p < 0.05) (n=6 replicates) **(J)** Representative frequency distribution plot of fluorescence-associated cell sorting of DNA content (DAPI) in control and importazole-treated iPSC-derived motor neurons (n=25,000 cells). **(K)** Percentage of cells in G1, S, and G2 phase of the cell cycle according to the distribution of DNA content across all cells in control and importazole conditions (n=4, 25,000 cells). **(L)** Timeline of iPSC-derived motor neuron generation treated with the importin-β antagonist IPZ at DIV10 and time-course of associated IPZ-washouts at days 1-4 with a 2-day recovery period following the washout. **(M)** Seven-day viability curves of IPZ-treated motor neurons with IPZ-washouts at days 1-4 or no washout, analyzed by two-way ANOVA with Tukey’s multiple-comparison test (p < 0.05) (n=3 replicates). **(N-Q)** qPCR quantification of cell-cycle genes (*CCND1, CCNB1, CCNE1,* and *CCNA2*) at indicated washout timepoints with 2-day recovery, shown as fold change relative to control, analyzed by one-way ANOVA with Tukey’s multiple-comparison test (p < 0.05) (n=3 replicates). Bar graphs represent mean ± standard deviation and significance values are indicated as * p<0.05, ** p<0.01, *** p<0.001. Scale bars: 20 μm (B). Replicates are independent differentiations to account for batch variability. Schematics created with BioRender.com. All data points and statistical comparisons are provided in the Source Data file.

To further delineate the dynamics of this response, we quantified the expression of key cell cycle regulators (*CCND1, CCNB1, CCNE1,* and *CCNA2*) by qPCR. Consistent with the immunostaining results, *CCND1, CCNB1,* and *CCNA2* showed strong upregulation between days 1 and 3, whereas *CCNE1* expression peaked at day 2 (Fig. 5D-E). These findings indicate that nuclear import disruption elicits a temporally coordinated yet transient wave of aberrant cell cycle activation, ultimately culminating in neuronal loss. We next examined neuronal integrity by assessing *STMN2* and its truncated transcript variant. Full-length *STMN2* expression decreased rapidly after treatment and remained suppressed throughout the time course, while truncated *STMN2* transcripts accumulated progressively over six days (Fig. 5H-I), consistent with impaired axonal maintenance.

Given the observed wave of aberrant cell-cycle activation, we next asked whether neurons progress beyond G1/S and initiate DNA replication following importazole treatment. To assess this, we performed fluorescence-associated cell sorting (FACS) to quantify DNA content in control and importazole-treated motor neurons at 2 days post-treatment. Control neurons showed a distribution of ∼79.8/18.6/1.6% (G1/S/G2) across 25,000 events, consistent with their post-mitotic state (Fig. 5J-K) and the high purity of our differentiation (Fig. 3B-C). In contrast, importazole-treated cells exhibited a shift to ∼71.9/26.0/3.0% (G1/S/G2), reflecting a ∼7.4% increase in S-phase cells. This redistribution indicates that a subset of neurons indeed attempts to initiate DNA replication in response to nuclear import stress. Notably, the absence of a corresponding accumulation in G2 suggests that these neurons fail to complete S-phase progression, consistent with an abortive cell-cycle response.

To determine whether transient inhibition of nuclear import is sufficient to trigger irreversible degeneration, we performed washout experiments in which importazole was removed daily from days 1-4 (Fig. 5L). Viability analyses revealed that washout after one day produced minimal neuronal loss, whereas removal at days 2 or 3 resulted in moderate but significant cell death (Fig. 5M). Washout at day 4 caused the highest level of neuronal death, comparable to continuous drug exposure over seven days (Fig. 5M). Follow-up qPCR performed 48 hours after each washout showed that early washout (days 1-3) prevented significant re-expression of cyclins, while washout at day 4 led to a pronounced induction of *CCND1, CCNE1,* and *CCNA2,* as well as an increase, but not significant increase in *CCNB1* (Fig. N-Q).

Together, these data indicate that importazole-induced nuclear import disruption initiates an early but reversible phase of cell-cycle activation that becomes irreversible after day 3, defining a critical temporal threshold for neuronal survival.

### DNA-replication events drive cell-cycle-associated neuronal cell death

To investigate whether DNA replication contributes to neuronal death following importazole-induced cell-cycle activation, we leveraged 5-fluoro-2′-deoxyuridine (FUDR), a thymidylate synthase inhibitor that blocks DNA synthesis (*44*). During differentiation, FUDR was routinely applied to eliminate proliferating cells and enrich for post-mitotic motor neurons. For this experiment, FUDR was removed three days prior to importazole treatment to allow neurons to recover. We then reintroduced FUDR simultaneously with importazole to assess the consequences of inhibiting DNA synthesis during nuclear import disruption (Fig. 6A).

**Figure 6:**
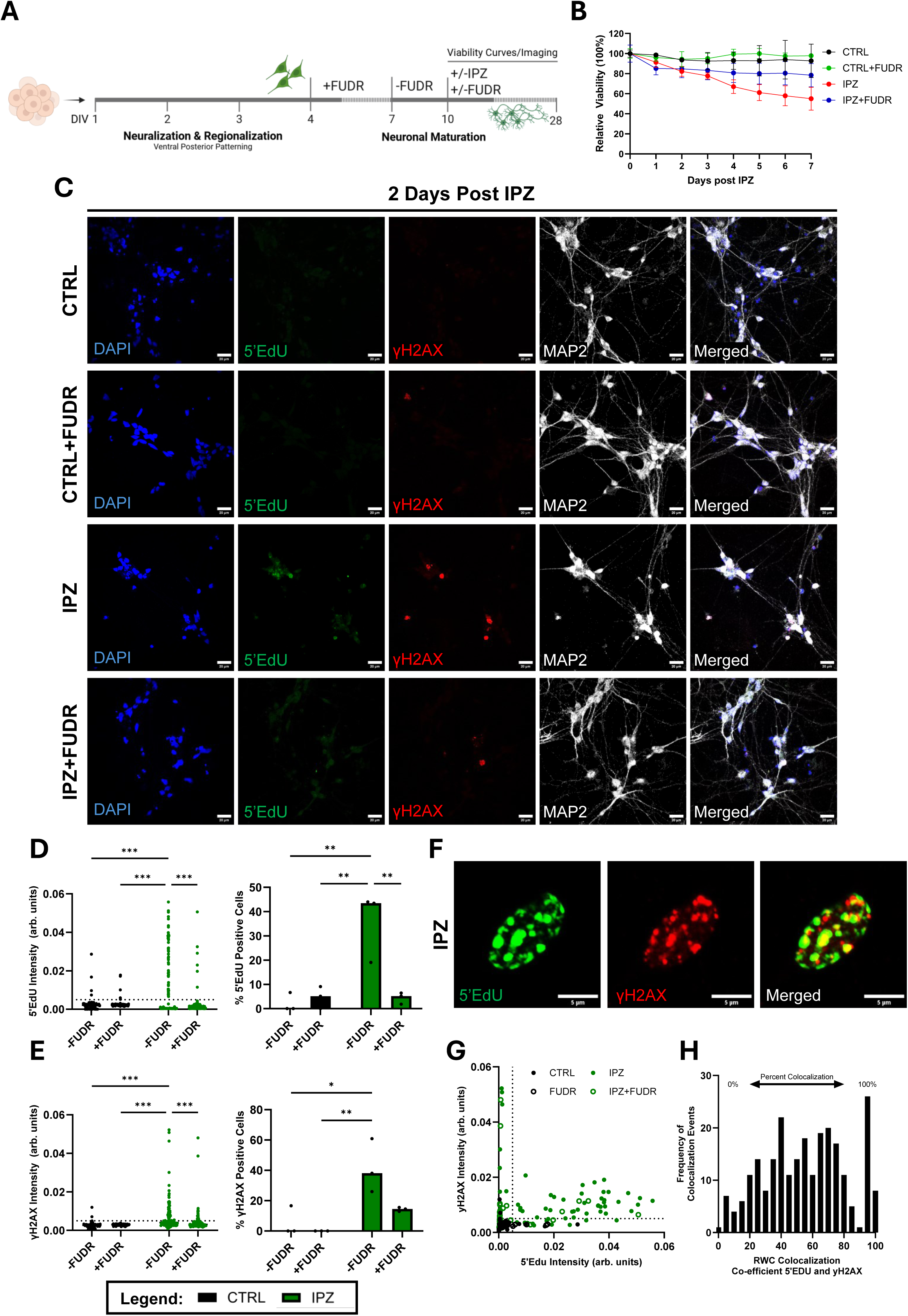
DNA-replication events drive cell-cycle-associated neuronal cell death. **(A)** Timeline of iPSC-derived motor neuron generation treated with DMSO control or IPZ treatment with or without FUDR at DIV10, and analyzed by viability assays or imaging at 2 days post IPZ. **(B)** Seven-day viability curves of motor neurons treated with DMSO control or IPZ treatment with or without FUDR, analyzed by two-way ANOVA with Tukey’s multiple-comparison test (p < 0.05) (n=3 replicates). **(C)** Representative immunostains of DAPI (blue), 5’EdU (green), γH2AX (red), and MAP2 (white) in iPSC-derived motor neurons 2 days after DMSO control or IPZ treatment, with or without FUDR. **(D)** Quantification of 5’EdU intensity and percentage of 5’EdU (mean intensity >0.005 threshold) positive motor neurons, analyzed by two-way ANOVA with Tukey’s multiple-comparison test (p < 0.05) (n=3 replicates). **(E)** Quantification of γH2AX intensity and percentage of γH2AX (mean intensity >0.01 threshold) positive motor neurons, analyzed by two-way ANOVA with Tukey’s multiple-comparison test (p < 0.05) (n=3 replicates). **(F)** Representative immunostains of 5’EdU (green) and γH2AX (red) in iPSC-derived motor neurons 2 days after IPZ treatment showing colocalization of signal. **(G)** Correlation plot of 5’EdU and γH2AX intensity (mean intensity >0.005 threshold) in iPSC-derived motor neurons 2 days after DMSO control or IPZ treatment, with or without FUDR from (C-E). **(H)** Frequency histogram showing RQC colocalization coefficient from 5’EdU and γH2AX immunostains from (F). Bar graphs represent mean ± standard deviation and significance values are indicated as * p<0.05, *** p<0.001. Scale bars: 20 μm (C) and 5 μm (F). Replicates are independent differentiations to account for batch variability. Schematics created with BioRender.com. All data points and statistical comparisons are provided in the Source Data file.

Viability analyses revealed that FUDR alone had no effect on neuronal survival, whereas importazole treatment led to significant cell death. Surprisingly, co-treatment with FUDR and importazole rescued the viability loss observed with importazole alone (Fig. 6B). To determine whether this effect was due to inhibition of DNA synthesis, we performed 5′-ethynyl-2′-deoxyuridine (EdU) incorporation assays after two days of treatment. Importazole induced EdU-positive nuclear foci in a subset of post-mitotic motor neurons, while co-treatment with FUDR significantly reduced EdU incorporation (Fig. 6C-D), consistent with inhibition of DNA synthesis.

We next assessed DNA damage using γH2AX. Importazole-treated neurons exhibited a robust increase in focal γH2AX, whereas FUDR co-treatment substantially reduced overall γH2AX burden (Fig. 6D-E). Importantly, within the importazole condition, the vast majority of EdU-positive neurons were also γH2AX-positive (Fig. 6F-G), and EdU foci showed near-complete colocalization with γH2AX (Fig. 6H), indicating that sites of aberrant DNA synthesis coincide with DNA damage signaling. Together, these findings support a model in which importazole triggers inappropriate DNA replication initiation that drives a replication-associated DNA damage response and neuronal death, and that blocking DNA synthesis with FUDR prevents this damage and improves survival. However, because γH2AX signal was not fully eliminated by FUDR co-treatment (Fig. 6E), importazole likely also induces a replication-independent DNA damage response, consistent with nuclear import disruption compromising genome maintenance pathways in neurons (*16, 45*).

To further assess the temporal sensitivity of this effect, FUDR was introduced at sequential time points following importazole exposure (days 0-4) (Fig. S7A). FUDR addition at day 0 prevented neuronal death, while introduction at days 1-3 exacerbated degeneration (Fig. S7B). In contrast, addition at day 4 had no significant effect compared to importazole alone (Fig. S7B). Altogether, these findings indicate that importazole-induced nuclear import disruption promotes aberrant DNA replication, leading to DNA damage and neuronal death, and that timely inhibition of DNA synthesis can mitigate these effects and restore neuronal survival.

### Inhibition CDK4/6 blocks DNA replication and offers neuroprotection

Having established that neurons tolerate early cell-cycle activation but succumb following DNA replication initiation, we next evaluated whether restricting G1/S checkpoint progression through CDK4/6 inhibition could prevent this lethal transition. To target CDK4/6, we utilized the FDA-approved inhibitors palbociclib and abemaciclib, both widely used in the treatment of breast cancer (36, 37). To control for off-target toxicity, we first performed dose–response viability assays in iPSC-derived motor neurons with and without importazole (IPZ) treatment (Fig. S8). Palbociclib increased survival of IPZ-treated neurons at concentrations between 1-10 nM, with minimal effects at lower doses and clear toxicity at higher concentrations (Fig. S8). Abemaciclib similarly enhanced survival at 1-5 nM, whereas concentrations above 10 nM produced significant toxicity (Fig. S8). Based on these dose–response curves, we selected 5 nM for both inhibitors for subsequent experiments, as this concentration provided maximal rescue with minimal independent toxicity (Fig. 7A-B). At this dose, neither palbociclib nor abemaciclib impaired long-term neuronal viability when administered alone.

**Figure 7:**
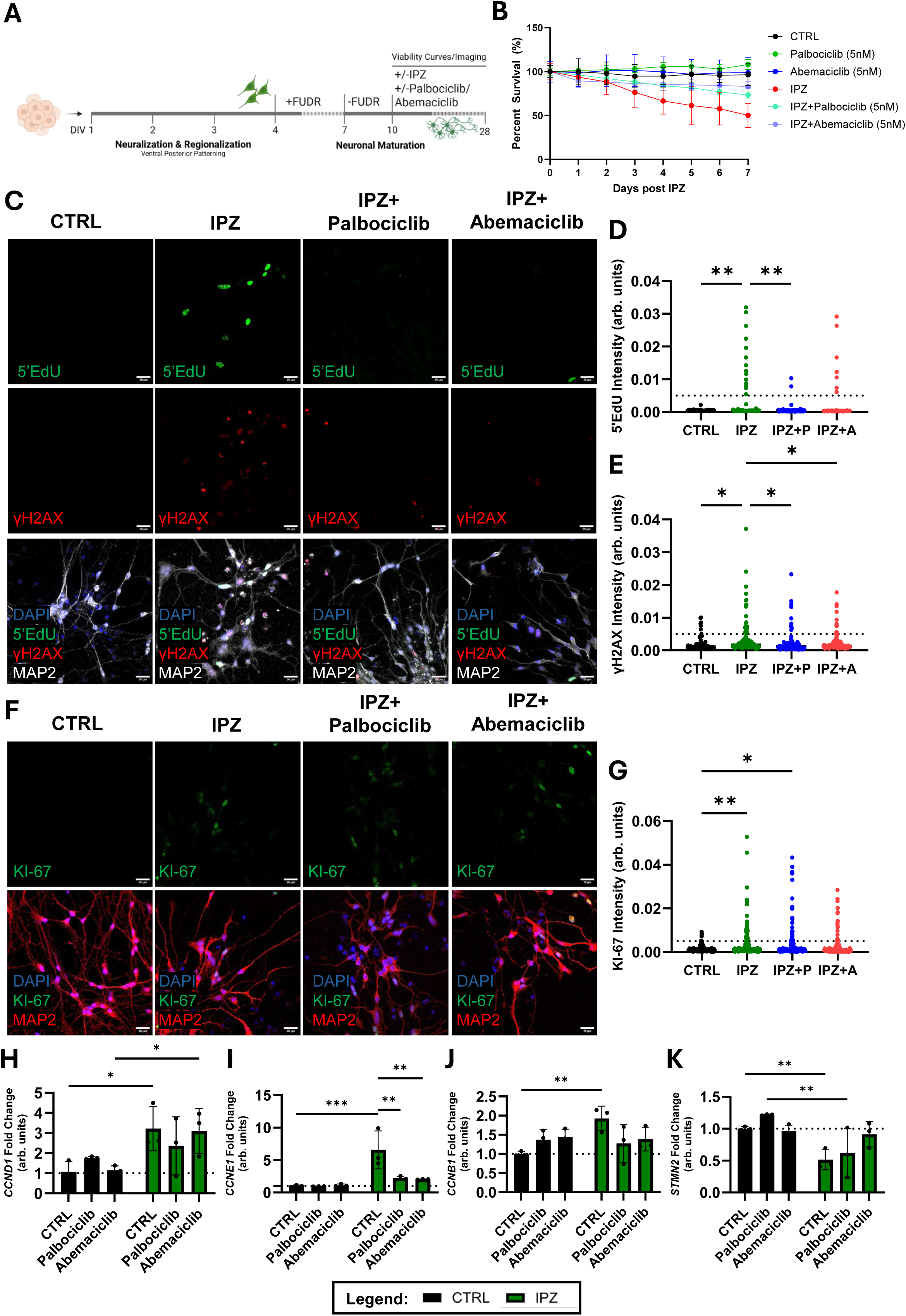
Inhibition CDK4/6 blocks S-phase entry and offers neuroprotection. **(A)** Timeline of iPSC-derived motor neuron generation treated with DMSO control or IPZ treatment with or without palbociclib or abemaciclib at DIV10, and analyzed by viability assays or imaging at 2 days post IPZ. **(B)** Seven-day viability curves of motor neurons treated with DMSO control or IPZ treatment with or without palbociclib or abemaciclib, analyzed by two-way ANOVA with Tukey’s multiple-comparison test (p < 0.05) (n=3 replicates). **(C)** Representative immunostains of DAPI (blue), 5’EdU (green), γH2AX (red), and MAP2 (white) in iPSC-derived motor neurons 2 days after DMSO control or IPZ treatment, with or without palbociclib or abemaciclib. **(D)** Quantification of 5’EdU intensity (mean intensity >0.005 threshold) in motor neurons, analyzed by one-way ANOVA with Holm-Šídák’s multiple comparisons test (p < 0.05) (n=3 replicates). **(E)** Quantification of γH2AX intensity (mean intensity >0.01 threshold) in motor neurons, analyzed by one-way ANOVA with Holm-Šídák’s multiple comparisons test (p < 0.05) (n=3 replicates). **(F)** Representative immunostains of DAPI (blue), Ki-67 (green), and MAP2 (red) in iPSC-derived motor neurons 2 days after DMSO control or IPZ treatment, with or without palbociclib or abemaciclib. **(G)** Quantification of Ki-67 intensity (mean intensity >0.005 threshold) in motor neurons, analyzed by one-way ANOVA with Holm-Šídák’s multiple comparisons test (p < 0.05) (n=3 replicates). (**H-K)** qPCR analysis of cell-cycle and disease-relevant genes (*CCND1*, *CCNB1*, *CCNE1*, and *STMN2*) in iPSC-derived motor neurons 2 days after DMSO control or IPZ treatment, with or without palbociclib or abemaciclib. Bar graphs represent mean ± standard deviation and significance values are indicated as * p<0.05, *** p<0.001. Scale bars: 20 μm (C) and (F). Replicates are independent differentiations to account for batch variability. Schematics created with BioRender.com. All data points and statistical comparisons are provided in the Source Data file.

When co-administered with IPZ, both CDK4/6 inhibitors conferred measurable neuroprotection over a 7-day time course; however, only abemaciclib achieved statistically significant rescue by day 7 (Day 7: IPZ vs. IPZ+palbociclib, p=0.1095; IPZ vs. IPZ+abemaciclib, p=0.0051) (Fig. 7B). To determine whether this protection resulted from inhibition of pathological DNA replication, we next assessed DNA synthesis using 5′-EdU incorporation and DNA damage using γH2AX immunostaining. Both palbociclib and abemaciclib markedly reduced EdU incorporation, confirming effective restriction of G1/S progression (Fig. 7C-D). In contrast, γH2AX levels were only modestly reduced (Fig. 7E), suggesting that aberrant DNA replication, rather than DNA damage alone, is the primary pathological driver downstream of nuclear import disruption.

Notably, CDK4/6 inhibition did not induce complete cell-cycle exit, as neither palbociclib nor abemaciclib reduced expression of the proliferation marker Ki-67 (Fig. 7F-G). qPCR analysis further supported this interpretation, revealing normalization of *CCNE1* and *CCNB1* expression without reduction of CCND1, consistent with arrest in G1 rather than terminal cell-cycle withdrawal (Fig. 7H-J). Finally, both inhibitors partially rescued expression of the neuronal integrity marker STMN2 (Fig. 7K), indicating that restricting pathological S-phase entry preserves aspects of neuronal health despite ongoing cellular stress.

Together, these findings demonstrate that CDK4/6 inhibition confers neuroprotection not by forcing cell-cycle exit, but by selectively preventing pathological S-phase entry and replication-associated DNA-damage downstream of nuclear import dysfunction.

## Discussion

Our findings demonstrate that patient stratification based on cell-cycle gene expression reveals meaningful biological heterogeneity within ALS, indicating that neuronal cell-cycle engagement is not uniformly detrimental but instead reflects divergent neuronal states. Rather than representing a uniform failure mode, our patient transcriptomic data and iPSC-derived neuron models reveal that cell-cycle re-entry emerges as a cellular stress response whose outcome depends on how far neurons progress past initiation. Early or partial activation, particularly stalled engagement during S-phase, predominantly leads to replication stress, DNA damage, and neuronal death, whereas neurons that successfully navigate beyond DNA replication and enter G2 correspond to patient subgroups with better functional outcomes. Thus, neuronal cell-cycle re-entry acts as a bifurcation point, where incomplete or aborted progression is harmful, but full passage through S-phase can be adaptive or neuroprotective. This framework helps explain patient heterogeneity and suggests that targeting the timing and extent of cell-cycle activation could be a key therapeutic strategy.

Decades of work have suggested that neuronal cell-cycle re-entry is uniformly pathological in ALS and other neurodegenerative diseases, largely based on observations of aberrant cyclin expression, DNA replication stress, and apoptotic markers in vulnerable neuronal populations (*1, 15, 28, 46–48*). However, our analysis of the AnswerALS cohort indicates a more nuanced landscape. Consistent with earlier reports of heterogeneous molecular phenotypes in ALS (*49–51*), we identify patient subgroups distinguished by differential engagement of cell-cycle programs. Notably, one cluster exhibits strong induction of G1 and S/G2 cyclins (*CCND1, CCNB1, CCNA2*) in the absence of *CCNE1* upregulation, a pattern reminiscent of the “abortive cell-cycle re-entry” described in prior neuropathological studies (*52*). Yet, when examined in the context of clinical progression, this activation is not uniformly detrimental. Instead, we observe a biphasic relationship in which moderate activation correlates with worse ALSFRS-R outcomes, whereas high expression of later-phase regulators corresponds to more favorable trajectories. These findings extend earlier hypotheses that neurons may engage cell-cycle machinery heterogeneously and suggest that the specific phase and extent of this activation, not simply the presence of cyclin expression, encode prognostic information (*6*).

Our mechanistic studies provide experimental grounding for this interpretation. Nuclear import defects have long been implicated in ALS pathogenesis, particularly in *C9orf72*-associated disease and models of TDP-43 pathology (*25, 53–56*), but their relationship to cell-cycle regulation has remained unclear. Using importazole to selectively inhibit importin-β, we recapitulated hallmark ALS features, including cytoplasmic pTDP-43 accumulation and widespread nuclear import failure, aligning with previous observations in patient tissue and genetic models. Importantly, this disruption provoked rapid induction of G1-, S-, and G2-associated cyclins, consistent with the transcriptional signatures observed in patient subgroups and reinforcing the idea that NCT dysfunction is an upstream driver of neuronal cell-cycle activation. SILAC-based spatial proteomics further revealed large-scale protein mislocalization affecting both cell-cycle assemblies and neurodegenerative pathways, supporting prior work demonstrating that loss of nuclear compartmentalization destabilizes proteostasis and signaling networks (*18, 20, 57*).

By integrating temporal analyses, we show that the neuronal response to nuclear import failure is dynamic and initially reversible, echoing earlier suggestions that neurons can tolerate brief periods of aberrant cell-cycle signaling. However, once neurons approach or enter S-phase, the trajectory becomes far more consequential. Consistent with studies demonstrating that DNA replication stress is a potent inducer of neuronal apoptosis (*46, 58*), we identify DNA-replication initiation, not early G1 activation, as the critical inflection point leading to DNA damage accumulation and cell death. Blocking DNA synthesis predominantly rescues viability, yet our transcriptomic patient data suggest that neurons capable of completing S-phase and reaching G2 may instead achieve a more stable, protective state. This observation aligns with recent proposals that some neurons adopt a pseudo-mitotic or tetraploid-like state under stress, providing short-term resilience (21, 48).

These mechanistic insights highlight multiple potential strategies for promoting neuronal survival following nuclear import disruption. Previous studies have demonstrated that modulation of DNA damage checkpoint signaling, including inhibition of kinases such as CHK2, can confer neuroprotection (*59, 60*), suggesting that relaxation of checkpoint enforcement may permit neurons to escape lethal arrest. Although these studies did not directly examine neuronal cell cycle re-entry, they point to checkpoint pathways as tractable therapeutic targets. In parallel, our findings identify CDK4/6 inhibition as an alternative strategy that intervenes earlier in the cell cycle, restraining S-phase entry without engaging DNA replication or downstream proliferative programs. This distinction may be particularly relevant in post-mitotic neurons, where uncoupling survival from replication is likely to be essential. While prior cancer-focused studies typically employ palbociclib and abemaciclib at micromolar concentrations (*52–54*), we demonstrate that ultra-low-dose CDK4/6 inhibition (5 nM) is sufficient to confer robust neuroprotection with minimal off-target toxicity. Together, these observations suggest that both checkpoint modulation and cell-cycle restriction represent promising, non-mutually exclusive therapeutic avenues, with CDK4/6 inhibition offering a mechanistically conservative approach tailored to the unique vulnerabilities of neurons (*61*).

Together, our patient and experimental data converge on a model in which disease-initiating insults, such as nuclear import defects, trigger a potentially adaptive reactivation of early cell-cycle programs that becomes pathological upon inappropriate replication initiation. This framework reconciles prior observations of neuronal cell-cycle markers in ALS with their inconsistent associations with disease severity. It further raises the possibility that therapeutic strategies aimed at stabilizing early G1 or preventing S-phase entry, such as low-dose CDK4/6 inhibition, may preserve neurons before irreversible replication stress occurs.

## Limitations of the Study

This study has several limitations. First, analyses using the AnswerALS cohort are inherently correlational and cannot establish causality between transcriptional signatures and clinical outcomes. Moreover, iPSC-derived motor neurons, although valuable for modeling patient-specific biology, may not fully recapitulate the maturation state, environmental context, or chronic stress experienced by in vivo motor neurons. Second, our use of importazole as a model for ALS-related NCT dysfunction provides a synchronized and temporally controlled system to dissect downstream pathways. Importazole induces rapid and relatively uniform disruption of Importin-β-mediated nuclear import, allowing alignment of cellular responses across time points. However, this acute, population-wide effect may not reflect the progressive and heterogeneous nature of NCT decline in human disease. In ALS, nuclear import defects likely accumulate gradually and only produce pathological outcomes once compensatory mechanisms are overwhelmed, suggesting a threshold-dependent process that is not fully modeled by acute chemical inhibition. Future work combining longitudinal patient datasets with chronic or low-grade NCT perturbations in genetic ALS models will be essential for determining whether the threshold-dependent cell-cycle phenotypes identified here extend across diverse disease subtypes and more accurately represent human disease progression.

## Methods

### AnswerALS Transcriptomic Analysis

Transcriptomic data for ALS patients and controls were obtained from the AnswerALS consortium database (*62*) (Neuromine, accessed July 7^th^, 2025). A total of 624 individuals (445 ALS, 161 healthy controls, 10 non-ALS motor neuron disease [MND] cases, and 8 asymptomatic ALS gene carriers) with available high-quality RNA-sequencing data were included in this study. Raw count matrices were normalized and analyzed using the *DESeq2* package (v1.40.2) in R (v4.3.1) to identify differentially expressed genes and to generate normalized expression values for downstream analyses.

Dimensionality reduction was performed using Uniform Manifold Approximation and Projection (UMAP) implemented through the *umap* package (v0.2.10.0) in R. K-means clustering (k = 10) was applied to the UMAP embeddings to identify transcriptionally distinct patient subgroups. For visualization and comparative analyses, gene expression values were z-scored across samples, and cluster assignments were used to stratify patients according to expression of selected cell cycle–related genes (*CCND1, CCNB1, CCNE1,* and *STMN2*).

Clinical metadata from, including ALS Functional Rating Scale–Revised (ALSFRS-R) scores (n=290) and survival times (n=227), were integrated with transcriptomic data to evaluate disease outcomes. Kaplan–Meier survival analyses and curve fitting were performed using GraphPad Prism (v10.0), with significance determined by the log-rank (Mantel–Cox) test. Longitudinal changes in ALSFRS-R scores were analyzed using linear mixed-effects models implemented in the *lme4* (v1.1-35.2) and *lmerTest* (v3.1-3) R packages. ALSFRS-R score was modeled as a function of time since the first visit, the grouping variable (either *cluster* or *cyclin expression level*), and their interaction term (*time x group*), with a random intercept included for each subject to account for repeated measures within individuals. The interaction term tested whether the rate of functional decline (slope) differed between groups. Significance of fixed effects was evaluated using Satterthwaite’s approximation for degrees of freedom. Post-hoc pairwise comparisons of the estimated time-dependent slopes were performed using the *emmeans* (v1.10.1) package, and *p*-values were adjusted for multiple testing using the false discovery rate (FDR) method. Groups or clusters exhibiting significant pairwise slope differences (adjusted *p* < 0.05) were annotated with asterisks in the corresponding trajectory plots or legends to indicate differential rates of disease progression.

### TetO-hNGN2*-*puro iPSC Generation

Doxycycline-inducible human Neurogenin-2 (*hNGN2*) iPSCs were generated as previously described (Cookson et al., 2022)(*63*) with minor modifications. iPSCs were maintained in StemFlex (Gibco) and grown on Matrigel (Corning) coated plates at 37℃ and 5% CO_2_ prior to transfection. On the day of the transfection (Day 0), iPSCs were dissociated with Accutase (Gibco) and 1.5 × 10^6^ cells were replated in a 6-well TC dish coated with Matrigel matrix and cultured in StemFlex supplemented with CEPT(*64*) (50nM Chroman 1 (MedChemExpress), 50µM Emricasan, 1X Polyamine Supplement (Sigma), 0.7 µM Trans-ISRIB (Tocris)). Cells were placed in an 37℃ 5% CO_2_ incubator for 2 hours. During the incubation time, a transfection mixture of 200µL OptiMEM (Gibco), 3µg DNA mix (1:3 EF1α Transposase:PB-TO-hNGN2-BFP plasmid ratio), 5µL Lipofectamine Stem (Invitrogen) was mixed and incubated separately for 30 minutes at room temperature. The transfection mixture was added dropwise to the iPSCs, and the plate was returned to the incubator. The following day, a positive nuclear BFP signal was confirmed, and cells were re-plated once 70%-90% confluency was achieved. Transfected iPSCs were selected with puromycin (8µg/mL) and clones were selected and amplified. TetO-*hNGN2-*puro iPSCs were frozen back at passage 22. Reagents and key resources are included in the Source Data file.

### *NGN2*-based iPSC-derived Spinal Motor Neuron Differentiation

TetO-hNGN2-puro iPSCs were maintained in StemFlex and grown on Matrigel coated plates at 37℃ and 5% CO_2._ On day 1, iPSCs were dissociated with Accutase and 1.5 × 10^6^ cells were replated in a 6-well TC dish coated with Matrigel matrix and grown in Induction Medium (DMEM/F-12 (Gibco), 1X N2 supplement (Gibco), 1X GlutaMAX (Gibco), 1X Non-essential amino acid (Gibco), 2µg/mL doxycycline hyclate (Sigma), 200nM LDN193189 (MedChemExpress), 10µM SB431542 (MedChemExpress), 1µM retinoic acid (MedChemExpress), and 1µM smoothened agonist (SAG, MedChemExpress)) until day 4 with daily media changes. On day 4, cells were dissociated with Accutase and replated 1:6 on Poly-L-Ornithine (Sigma) coated plates in neuronal supportive media (1:1 DMEM/F-12:BrainPhys (STEMCELL Technologies), 1X B27 Plus (Gibco),1X GlutaMAX, 10ng/mL BDNF, 10ng/mL GDNF, 10ng/mL NT-3, 1µM SAG, 1µM RA, 1µg/mL Laminin (R&D Systems), 1µM 5’-Fluoro-2’-deoxyuridine (FUDR, Sigma)). On day 7 and onwards, cells were cultured in neuronal maturation media (BrainPhys, 1X B27 Plus, 1X GlutaMAX, 10ng/mL BDNF, 10ng/mL GDNF, 10ng/mL NT-3, 1µM SAG, 1µM RA, 1µg/mL Laminin). Reagents and key resources are included in the Source Data file.

### Importazole and FUDR treatments

iPSC-derived spinal motor neurons were matured to day in vitro (DIV) 10 before adding either DMSO control (0.1% DMSO), 10µM importazole, and/or 1µM FUDR. Treatments were kept for the time courses indicated for each experiment.

### SILAC Labeling and Fractionation

iPSCs were differentiated as described (n=3 independent differentiations), with the modification of using DMEM/F-12 for SILAC (Thermo Scientific) supplemented with ‘heavy’ lysine and arginine isotopes (13C6, 15N2-Lys and 13C5,15N4-Arg), or ‘light’ isotopes (12C and 14N Lys and Arg) (Thermo Scientific). To replace BrainPhys, we utilized DMEM for SILAC (Thermo Scientific) supplemented with ‘heavy’ or ‘light’ lysine and arginine isotopes and 1X NEAA (Gibco), 0.227mM Sodium Pyruvate (Sigma), 5nM Vitamin B-12 (Sigma), 0.674µM Zinc Sulfate, and 25mM D-glucose.

2 million cells cultured in ‘heavy’ and ‘light’ media were separately lysed using a hypotonic lysis buffer (10mM Tris-Cl pH 7.4, 10mM NaCl, 2mM MgCl2, and 0.5% NP-40) for 10mins at 4℃. Cells were centrifuged at 500 x g for 5 mins and supernatant was collected as the cytoplasmic fraction. Pellets were washed twice with hypotonic lysis buffer. Cells were then resuspended in hypotonic lysis buffer nuclei and were triturated with a 20 ½g needle for 10 passes. Cells were spun down at 17,000 x g for 5mins and supernatant was collected. ‘Light’ cytoplasmic fractions and ‘heavy’ nuclear fractions were pooled together for LC-MS/MS analysis. Reagents and key resources are included in the Source Data file.

### LC-MS/MS and Proteomic Quantification

SILAC samples were processed by the Stony Brook University Proteomic Core. In brief, samples were adjusted to 5% SDS, 100mM triethyl ammonium bicarbonate (TEAB), 10mM dithiothreitol (DTT) in 100µl and incubated at 55℃ for 30 min. Proteins were alkylated in 25mM iodoacetamide (IAc) at RT, for 30 min in the dark. Proteins were precipitated by addition 10µl of 12% phosphoric acid, followed by 700µl of S-Trap bind/wash buffer (90% methanol/ 50 mM TEAB) to produce a micro precipitate. Samples were then loaded on an S-Trap mini cartridge (cat. # K02-mini-10 Protifi), washed four times with 90% methanol, 100mM TEAB, each followed by centrifugation at 4000 x g for 1 minute. The samples were digested with trypsin (20µg) in 50 mM TEAB in a humified incubator overnight at 37℃. Peptides were eluted by sequential addition of 80µl 50 mM TEAB, 0.2% formic acid, and 50% acetonitrile, 0.2% formic acid, each followed by centrifugation at 4000 x g for 1 minute. The samples were then dried by SpeedVac and resuspended in 0.1% formic acid (FA) and peptides analyzed by C18 reverse phase LC-MS/MS. HPLC C18 columns were prepared using a P-2000 CO_2_ laser puller (Sutter Instruments) and silica tubing (100µm ID x 20 cm) and were self-packed with 3u Reprosil resin. Peptides typically were separated using a flow rate of 300 nl/minute, and a gradient elution step changing from 0.1% formic acid to 40% acetonitrile (ACN) over 90 minutes, followed 90% ACN wash and re-equilibration steps.

Peptide identification and quantitation was performed using an orbital trap (Q-Exactive HF; ThermoFisher) instrument followed by protein database searching using Proteome Discoverer 2.4. Replicate samples were analyzed, using two different HPLC gradient profiles (0-30% ACN over 90’ and 0-40% ACN over 90’). Electrospray ionization is achieved using spray voltage of ∼2.3 kV. Information-dependent MS acquisitions are made using a survey scan covering m/z 375 – 1400 at 60,000 resolution, followed by ‘top 20’ consecutive second product ion scans at 15,000 resolution. AGC targets for MS and MS/MS are 1X10^6^ and 2X10^5^, maximum IT of 100ms and 50ms, an MS/MS loop size of 20 and dynamic exclusion for 20s. Mass resolution cutoffs for MS and MS/MS are 10ppm and 0.1 Da respectively. Data files were acquired with Xcalibur. Peptide alignments and quantitation were performed using Proteome Discoverer v3.1 software (ThermoFisher). Protein false discovery rates experiments are binned at 0.01 and 0.05 FDR. Peptide and PSM FDR cutoffs are typically set to 0.05. Two missed tryptic cleavages were allowed and modifications considered included static cysteine derivatization, and variable deamidation (NQ), water loss (ST), and oxidation (M). SILAC ‘heavy’ (13C6, 15N2-Lys and 13C5,15N4-Arg), or ‘light’ (12C and 14N Lys and Arg) were used. Pairwise peptide heavy/light ratios allow abundance calculations and t-test. The human reviewed UniProt dataset (50227 entries) are used for data alignment. Matched peptide-based label free quantitation and p-values were calculated by Benjamini-Hochberg correction for FDR.

### RNA Isolation and Quantitative PCR

Total RNA was isolated from iPSC-derived spinal motor neurons (n=6 independent differentiations) at the indicated time points using the PureLink RNA Mini Kit (Invitrogen) according to the manufacturer’s protocol. RNA concentration and purity were assessed using a NanoDrop spectrophotometer (Thermo Fisher Scientific), and samples were snap-frozen at −80°C until use. For cDNA synthesis, 200 ng of total RNA was reverse transcribed using SuperScript IV Reverse Transcriptase (Thermo Fisher Scientific) in a 20 μL reaction volume, which was subsequently diluted to 40 μL with nuclease-free water to yield a working concentration of 5 ng/μL. Quantitative real-time PCR (qPCR) was performed using SYBR Green PCR Master Mix (Applied Biosystems) on a QuantStudio 3 Real-Time PCR System (Applied Biosystems). Reaction specificity was confirmed by melt curve analysis. All control and experimental samples for a given target gene were analyzed on the same qPCR plate and run in triplicate. Oligonucleotide sequences and key resources are included in the Source Data file.

### Cell Viability Assays

Cell viability was assessed using the CCK-8 assay (GlpBio) according to the manufacturer’s instructions. iPSC-derived spinal motor neurons were plated in 96-well plates and maintained until DIV 10. Viability was measured for each condition, with DIV 10 (day 0) serving as the reference point. For each biological replicate (n=3 independent differentiations), values from three technical wells were averaged to control for seeding variability.

### Nuclear Import Reporter Assay

Nuclear import was assessed using a lentiviral construct overexpressing a GFP with a canonical nuclear localization sequence (3x SV-40 NLS) under a human synapsin promoter (pLenti-hSyn-nucGFP)(*38*). 2^nd^ generation lentivirus was produced by transfection of 293 cells (Human Embryonic Kidney Cell line) with the transgene (pLenti-hSyn-nucGFP), packaging plasmid (psPAX2) and envelope plasmid (pMD2.G). Differentiated spinal motor neurons were transduced with the virus for 72 hours prior to importazole or DMSO control treatment for 48 hours. pLenti-hSyn-nucGFP was a gift from Lorenz Studer (Addgene plasmid # 140190; http://n2t.net/addgene:140190; RRID:Addgene_140190). psPAX2 and pMD2.G was a gift from Didier Trono (Addgene plasmid # 12260; http://n2t.net/addgene:12260; RRID:Addgene_12260).

### DNA synthesis (5’EdU incorporation) assay

DNA synthesis was assessed through 5′-ethynyl-2′-deoxyuridine (EdU) incorporation. iPSC-derived spinal motor neurons were incubated with 10 µM EdU (Sigma) for 1-2 hours prior to fixation at the indicated time points. Cells were fixed with 4% paraformaldehyde for 15 minutes at room temperature, permeabilized with 0.1% Triton X-100 for 15 minutes, and incubated with a Click-iT reaction cocktail (1mM Copper Sulfate [Sigma], 5mM Ascorbic Acid [Sigma], 50 mM Tri-HCl [Sigma], 150 mM NaCl [Sigma], 4 μM Azide-488 [Sigma]) to fluorescently label incorporated EdU. Spinal motor neurons were labeled with MAP2 and nuclei were counterstained with Hoechst 33342, and coverslips were mounted with ProLong Glass Antifade reagent. Images were acquired using the FV3000 (Olympus) confocal fluorescence microscope under identical acquisition settings. For each condition, the intensity of EdU-positive nuclei was quantified across n ≥ 3 fields of view using ImageJ and CellProfiler. Antibodies and key resources are included in the Source Data file.

### Fluorescence Associated Cell Sorting (FACS) Cell Cycle Analysis

iPSC-derived spinal motor neurons were washed once with 1× PBS and dissociated using Accutase. Cells were pelleted by centrifugation at 500 × g for 5 min, resuspended in 1 mL of ice-cold 70% ethanol, and fixed for at least 2 h with gentle inversion. Fixed cells were washed once with 1× PBS, centrifuged at 500 × g for 5 min, and resuspended in 500 μL of DAPI staining buffer (PBS containing 0.1% Triton X-100). Cells were incubated for 30 min at room temperature in the dark. Cell-cycle distribution was analyzed by flow cytometry with excitation at 340–380 nm and detection of DAPI fluorescence to discriminate G1, S, and G2/M phases. Cell-cycle phase discrimination and quantification were performed using ModFit LT.

### Palbociclib and Abemaciclib Treatments

iPSC-derived spinal motor neurons were matured to day in vitro (DIV) 10 before adding either DMSO control (0.1% DMSO), 10µM importazole, and/or 1µM Palbociclib, or 200nM Abemaciclib. Drug treatment concentrations were determined from drug viability assays. Treatments were kept for the time courses indicated for each experiment.

### Immunostaining

iPSC-derived spinal motor neurons were fixed with 4% paraformaldehyde for 15 minutes at room temperature and permeabilized with 0.1% Triton X-100 for 15 minutes. Cells were blocked for 1 hour at room temperature in blocking buffer containing 5% normal goat serum (NGS; Gibco) and 0.1% Tween-20 in PBS. Samples were incubated overnight at 4°C with the respective primary antibody diluted in blocking buffer. The following day, cells were washed four times for 5 minutes each with PBS containing 0.1% Tween-20 and incubated with fluorescent secondary antibodies in blocking buffer for 2 hours at room temperature in the dark. After four additional washes, coverslips were mounted with ProLong Glass Antifade reagent (Thermo Fisher). Images were acquired on an FV3000 confocal microscope (Olympus) under identical acquisition settings. Post-processing and quantitative image analysis were performed using ImageJ and CellProfiler. For each condition (n=3 independent differentiations), immunostaining quantifications were averaged across n ≥ 3 fields of view. Antibodies and key resources are included in the Source Data file.

### Figure design and visualization

Graphical illustrations were designed and prepared with Biorender.com and Adobe Illustrator. Plots were generated with R and GraphPad Prism and assembled in Adobe Illustrator.

### Statistical Analyses

Independent iPSC differentiations were treated as biological replicates, with each differentiation performed separately from thawing through maturation to capture biological variability inherent to the differentiation process. All statistical analyses were performed using GraphPad Prism (v10.0) and R (v4.3.1) unless otherwise specified. For transcriptomic analyses, normalization and differential expression were conducted using the DESeq2 package, and adjusted *p*-values were calculated using the Benjamini–Hochberg false-discovery-rate (FDR) method, with FDR < 0.05 considered significant. Kaplan–Meier survival analyses were generated in GraphPad Prism, and significance was determined by the log-rank (Mantel–Cox) test. For correlation analyses, Pearson’s correlation coefficients were calculated. Linear mixed-effects modeling, post-hoc slope comparisons, and model-based visualizations used in longitudinal analyses are described in detail under *AnswerALS transcriptomic analyses*.

For cell-based experiments, data are presented as mean ± standard deviation of the mean (SD) unless otherwise indicated. Comparisons between two groups were made using unpaired two-tailed Student’s *t*-tests with Bonferroni’s correction, and multiple-group comparisons were evaluated by one-way or two-way ANOVA followed by Tukey’s or S ida k’s post hoc tests as appropriate.

For imaging-based quantifications (Ki-67, EdU, γH2AX, and pTDP-43), 3 independent differentiations were analyzed per condition, with ≥3 fields of view per replicate. Proteomic *p*-values were adjusted for multiple comparisons using the Benjamini–Hochberg correction at an FDR< 0.05. Statistical significance for all analyses was defined as *p*<0.05.

## Data Availability

Data used in the preparation of this article were obtained from the AnswerALS Data Portal **(AALS-01184)**. For up-to-date information on the study, visit https://dataportal.answerals.org. All data generated or analyzed during this study are included in the accompanying Source Data file. This file contains processed proteomic and quantitative datasets, as well as underlying numerical values for all main and supplementary figures.

## Code Availability

All R scripts used for transcriptomic and statistical analyses in this study are available on GitHub at [https://github.com/jonathanplessis-belair/Neuronal-CCR-in-ALS]. The repository includes code for UMAP dimensionality reduction, DESeq2-based differential expression, ALSFRS-R trajectories, and survival analyses.

## Acknowledgements

A special thanks to John Haley and the Stony Brook University Proteomic Core for the SILAC LC/MS/MS analyses and to Todd Rueb at the Flow Cytometry Research Core for the Cell Cycle Analyses. Clinical data and biosamples used in the preparation of this article were obtained from the Answer ALS Foundation Program, ‘Answer ALS’. For up-to-date information on the program, <u>visit</u> https://www.answerals.org.

## Contributions

Study was conceived and designed by J.P.B and R.B.S. All experiments and analyses were performed by J.P.B. with supervisorial input from R.B.S. The manuscript draft was written by J.P.B. with input from R.B.S. Interpretation of data and contributions to the write-up was provided by J.P.B. and R.B.S.

