## Supplementary information for "Neuronal Cell-Cycle Re-entry Defines Divergent Outcomes Through Replication-Dependent DNA Damage in ALS"

| Cohort | Total | Clinical Mutation | | Sex | | Race | | | | Site of Onset | | | | | Average Age at Symptom Onset | Average Age at Death |
|---|---|---|---|---|---|---|---|---|---|---|---|---|---|---|---|---|
| - | - | Sporadic | Familial | Male | Female | White | Black/ African American | Asian | Other/NA | Limb | Axial | Bulbar | Multiple | Other/NA | - | - |
| **ALS** | 445 | 394 | 51 | 282 | 163 | 411 | 20 | 7 | 7 | 308 | 7 | 87 | 36 | 7 | 56.2 | 62.4 |
| **Asymptomatic ALS Gene carrier** | 8 | 0 | 8 | 3 | 5 | 8 | 0 | 0 | 0 | - | - | - | - | - | - | - |
| **Non-ALS MND** | 10 | - | - | 7 | 3 | 10 | 0 | 0 | 0 | 4 | 1 | 4 | 0 | 1 | 53.1 | 75.5 |
| **Healthy Control** | 161 | - | - | 75 | 86 | 62 | 3 | 3 | 93 | - | - | - | - | - | - | - |

**Table S1: Patient Cohort Demographic Data**

Supplementary figure is related to Figure 1.

| Cluster | Total | Clinical Mutation | | | Sex | | Race | | | | Site of Onset | | | | | Average Age at Symptom Onset | Average Age at Death |
|---|---|---|---|---|---|---|---|---|---|---|---|---|---|---|---|---|---|
| - | - | Sporadic | Familial | Healthy/ Other | Male | Female | White | Black/ African American | Asian | Other/NA | Limb | Axial | Bulbar | Multiple | Other/NA | - | - |
| **1** | 60 | 32 | 7 | 21 | 3 | 57 | 41 | 4 | 0 | 15 | 22 | 1 | 10 | 4 | 23 | 60.1 | 63.7 |
| **2** | 84 | 44 | 8 | 32 | 3 | 81 | 60 | 5 | 4 | 15 | 34 | 0 | 13 | 4 | 33 | 58.1 | 62.5 |
| **3** | 52 | 29 | 4 | 19 | 8 | 44 | 48 | 1 | 0 | 3 | 19 | 0 | 13 | 1 | 19 | 58.5 | 67.9 |
| **4** | 75 | 49 | 8 | 18 | 63 | 12 | 63 | 0 | 1 | 11 | 44 | 0 | 4 | 4 | 23 | 52.9 | 59.8 |
| **5** | 54 | 40 | 2 | 12 | 33 | 21 | 48 | 0 | 2 | 3 | 31 | 1 | 8 | 3 | 11 | 54.8 | 64.4 |
| **6** | 73 | 51 | 6 | 16 | 72 | 1 | 55 | 3 | 2 | 12 | 38 | 1 | 7 | 10 | 17 | 56.1 | 63.0 |
| **7** | 69 | 45 | 8 | 16 | 47 | 22 | 62 | 1 | 0 | 6 | 31 | 4 | 15 | 4 | 15 | 57.5 | 63.6 |
| **8** | 58 | 37 | 6 | 15 | 50 | 8 | 44 | 2 | 0 | 12 | 31 | 1 | 9 | 3 | 14 | 54.7 | 59.8 |
| **9** | 42 | 30 | 4 | 8 | 41 | 1 | 33 | 1 | 0 | 8 | 26 | 0 | 7 | 1 | 8 | 56.6 | 63.3 |
| **10** | 57 | 36 | 6 | 15 | 47 | 10 | 37 | 5 | 1 | 14 | 36 | 0 | 5 | 2 | 14 | 53.3 | 58.9 |

**Table S2: Patient Cluster Demographic Data**

Supplementary figure is related to Figure 1.

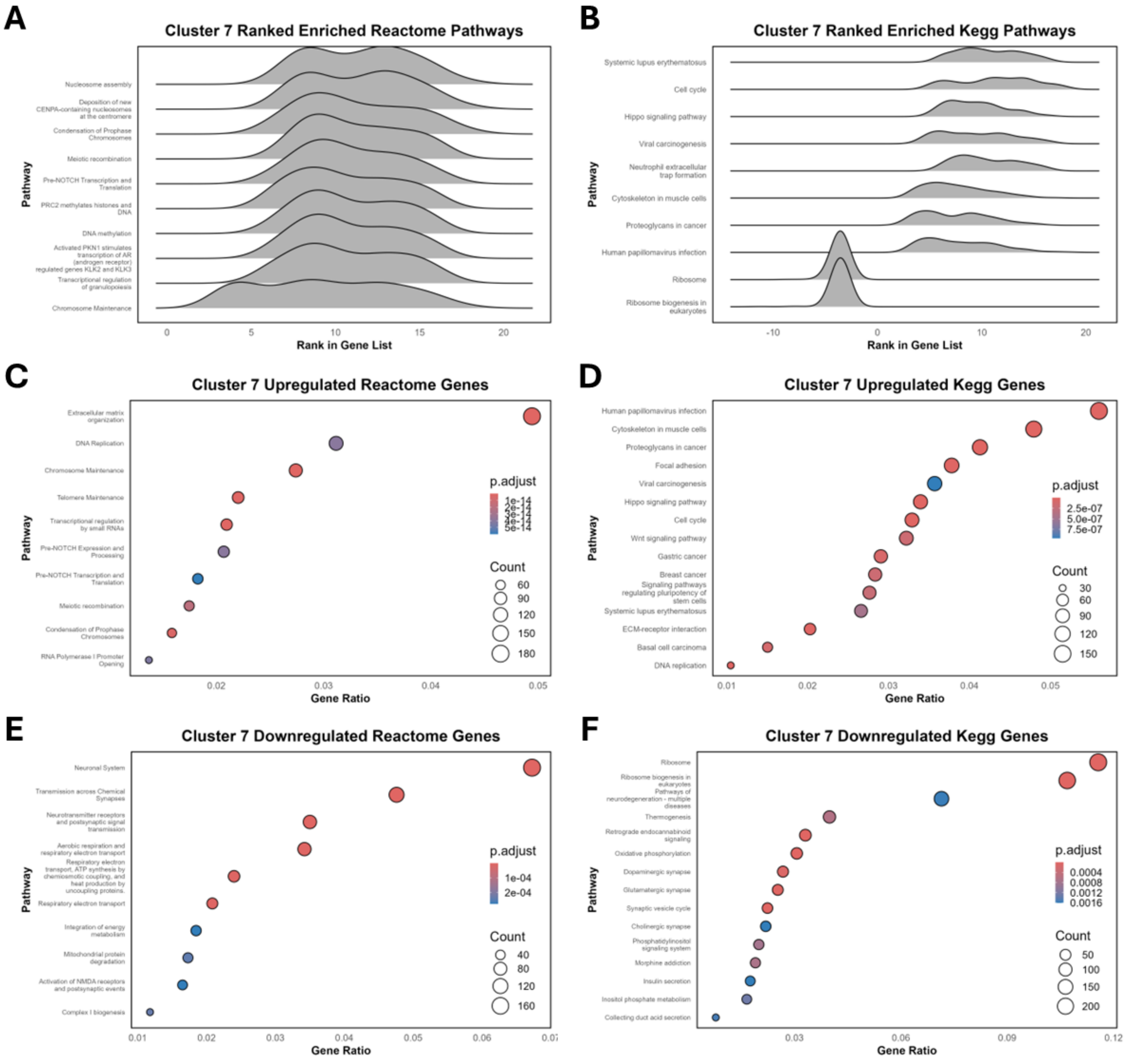


**Figure S1: Gene set enrichment analysis of cluster 7 from AnswerALS cohort. (A–B)** Ridge plots showing Reactome and KEGG pathway enrichment ranked by Wald statistic for cluster 7 of the AnswerALS cohort. **(C–F)** Dot plots showing Reactome and KEGG pathway enrichment ranked by Bonferroni-adjusted p-values (*p.adjust*) for cluster 7 of the AnswerALS cohort. Pathways are separated into upregulated (log2FC > 1, *p.adjust* < 0.05) and downregulated (log2FC < –1, *p.adjust* < 0.05) gene sets. Supplementary figure is related to Figure 1.

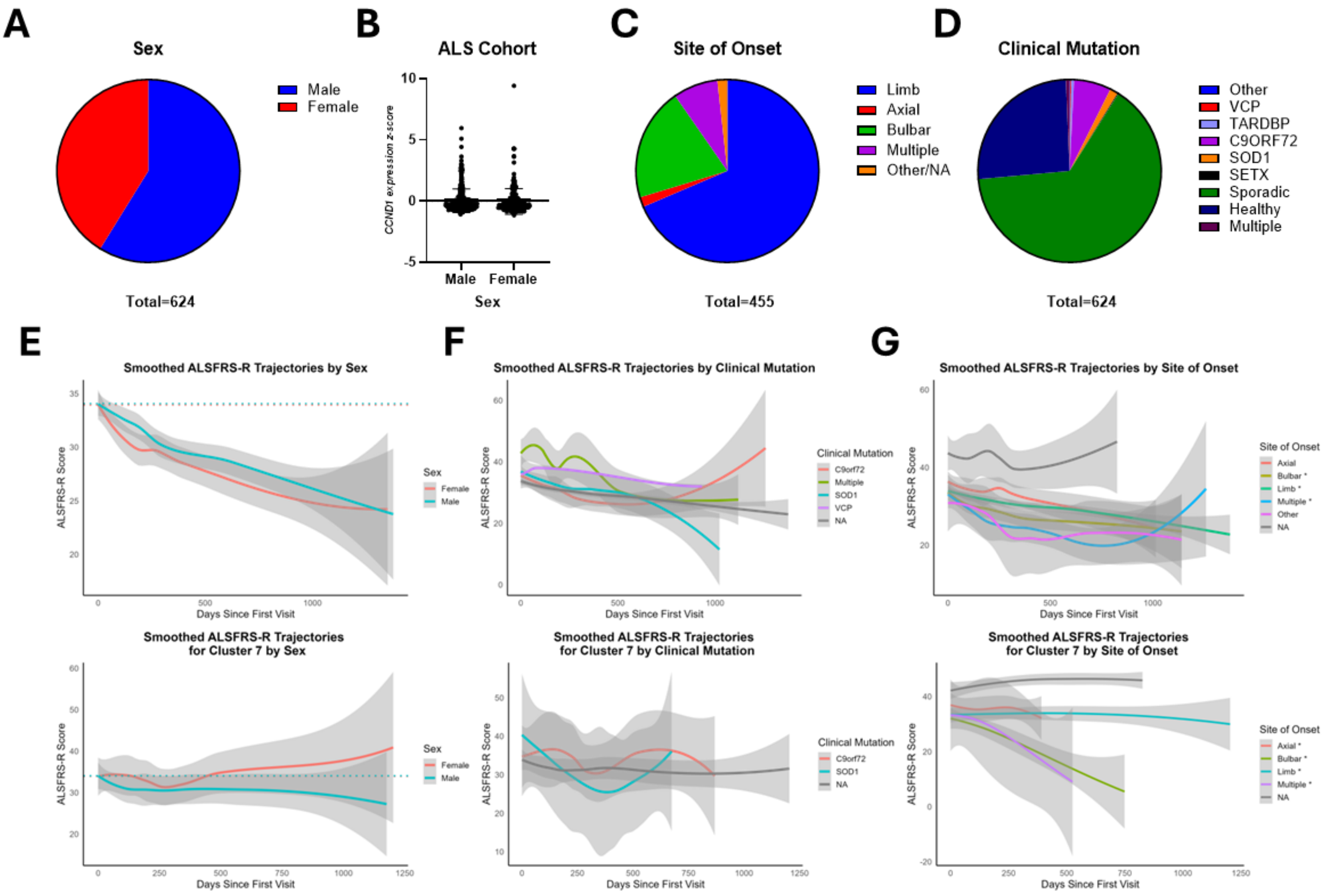


**Figure S2: AnswerALS patient trajectories by sex, clinical mutation, and site of onset. (A)** Pie chart showing the proportion of males and females in the AnswerALS cohort. **(B)** Bar graph displaying pairwise comparisons of CCND1 expression (z-score) between males and females in the AnswerALS cohort. **(C)** Pie chart illustrating the distribution of ALS patients by site of onset, including limb, axial, bulbar, multiple, and other/NA categories. **(D)** Pie chart illustrating the distribution of ALS patients by clinical mutation, including VCP, TARDBP, C9ORF72, SOD1, SETX, sporadic (NA), healthy, multiple, or other. **(E-G)** Smoothed ALSFRS-R trajectories across ALS patients (n=290) and cluster 7 (n=35) by sex, site of onset, and clinical mutation, with group differences evaluated using linear mixed-effects models. Supplementary figure is related to Figure 2.

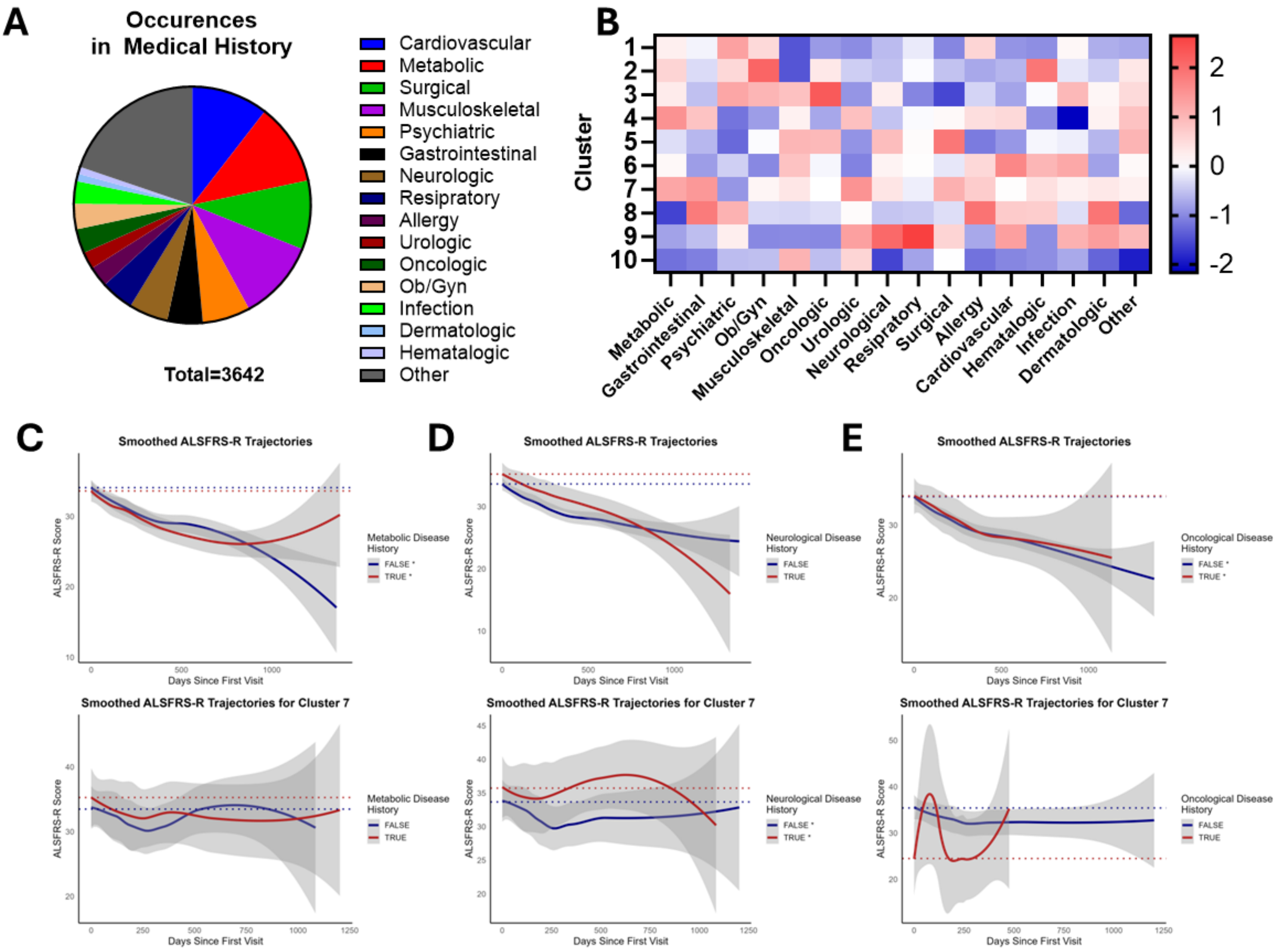


**Figure S3: AnswerALS patient medical history and comorbidities predict patient ALSFRS-R trajectories (A)** Pie chart showing the distribution of patients with the indicated medical history categories in the AnswerALS cohort. **(B)** Z-score heatmap illustrating the distribution of medical history categories across patient clusters. **(C–E)** Smoothed ALSFRS-R trajectories for all ALS patients (n = 290) and for cluster 7 (n = 35), stratified by medical history category. Group differences were evaluated using linear mixed-effects models. Supplementary figure is related to Figure 2.

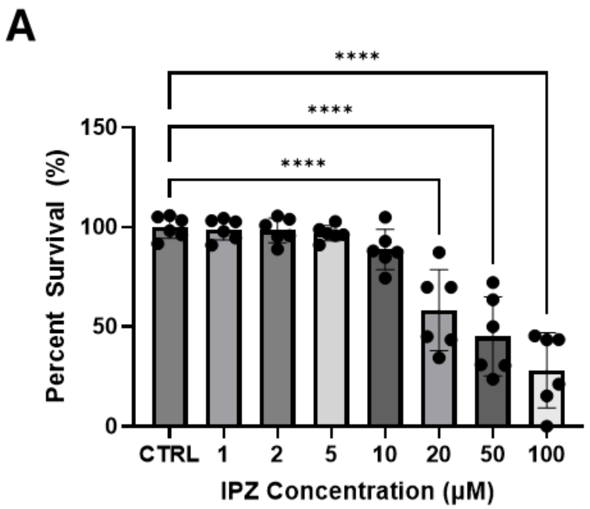


**Figure S4: Importazole dose-response curve (A)** Dose-response curve with increasing concentrations of importazole (IPZ) from 1µM to 100µM, analyzed by one-way ANOVA with Dunnett's multiple comparisons test ($p < 0.05$) (n=3 replicates). Bar graphs represent mean ± standard deviation and significance values are indicated as * p<0.05, *** p<0.001. Replicates are independent differentiations to account for batch variability. All data points and statistical comparisons are provided in the Source Data file.

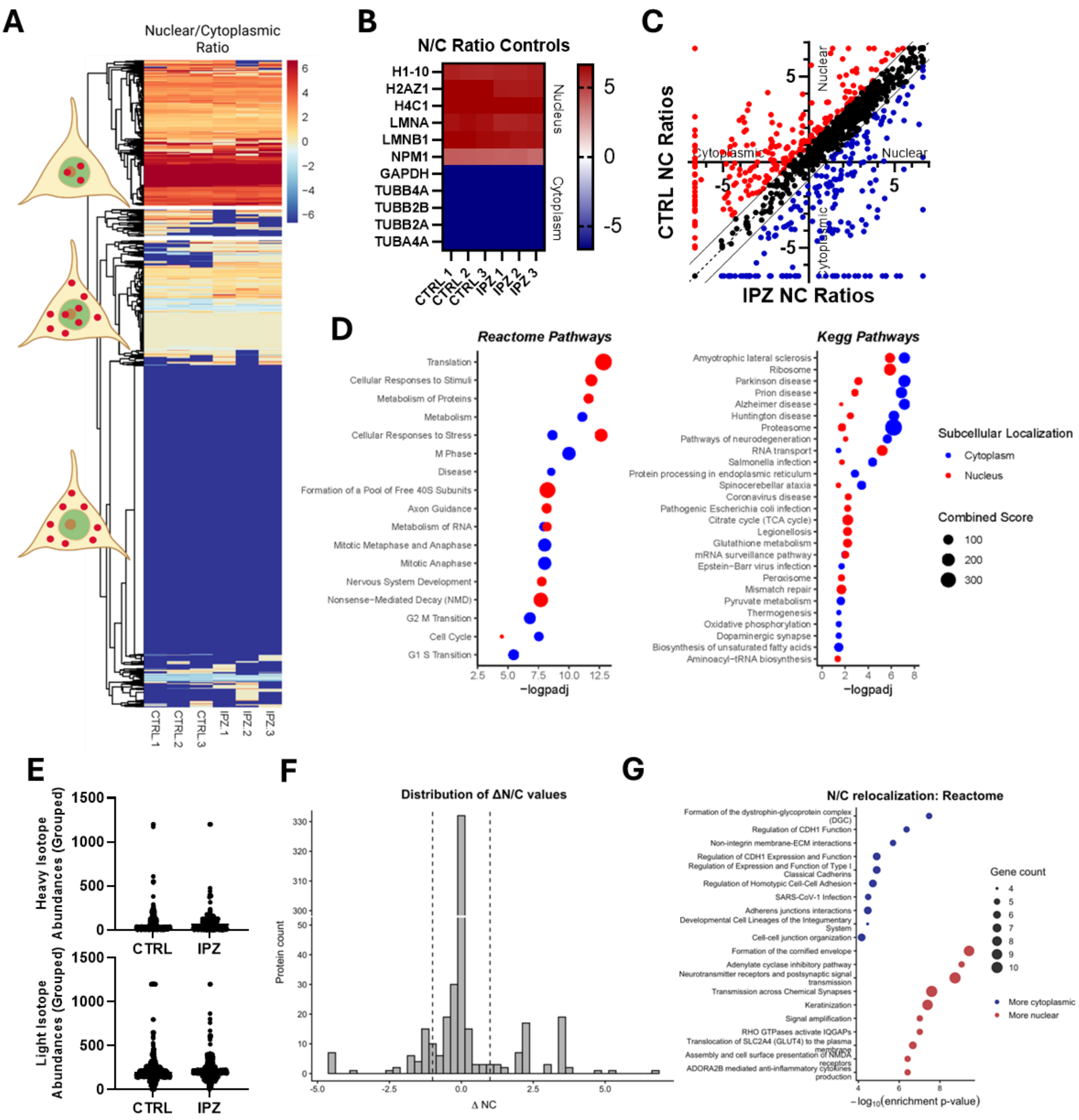

**Figure S5: SILAC-based spatial proteomics in a neuronal cell line and iPSC-derived motor neurons (A)** Heatmap depicting $\log_2$ fold-change nuclear-to-cytoplasmic (N/C) ratios of detected peptides in the SK-N-MC neuronal cell line, with nuclear enrichment shown in red, cytoplasmic enrichment in blue, and evenly distributed proteins in white (n = 3 biological replicates; Source Data). **(B)** Heatmap of known nuclear- and cytoplasmic-localized proteins used as subcellular fractionation controls. **(C)** Scatter plot showing the distribution of $\log_2$ fold-change N/C ratios under control (CTRL) and importazole (IPZ) conditions. **(D)** Reactome and KEGG pathway enrichment analysis of nuclear- and cytoplasmic-enriched differentially localized proteins following importazole treatment in SK-N-MC cells. **(E)** Pooled abundance of ‘heavy’ (nuclear) and ‘light’ (cytoplasmic) isotopes in CTRL and IPZ conditions for iPSC-derived motor neurons corresponding to Figure 4. **(F)** Distribution plot of differential N/C ratios (IPZ – CTRL) in iPSC-derived motor neurons, highlighting proteins with increased nuclear or cytoplasmic localization following importazole treatment. **(G)** Reactome pathway enrichment analysis of nuclear- and cytoplasmic-enriched differentially localized proteins following importazole treatment in iPSC-derived motor neurons.

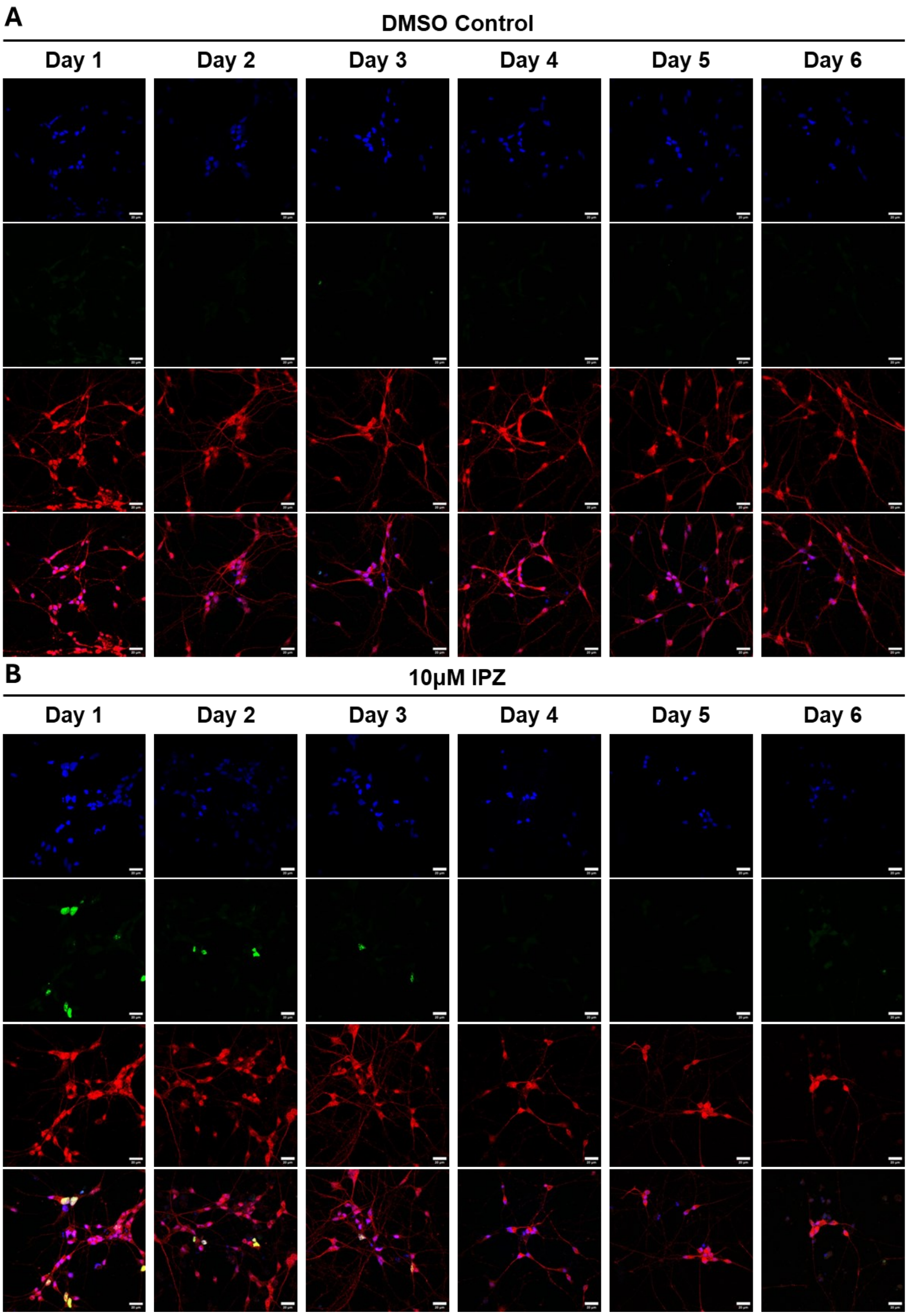


**Figure S6: Time-course of KI-67 expression in iPSC-derived motor neurons post IPZ treatment. (A)** Representative immunostains of KI67 (green), MAP2(red) in iPSC-derived motor neurons at DIV10. DAPI (blue) is a nuclear counterstain in the merged image (n=3 replicates). Supplementary figure is related to Figure 5.

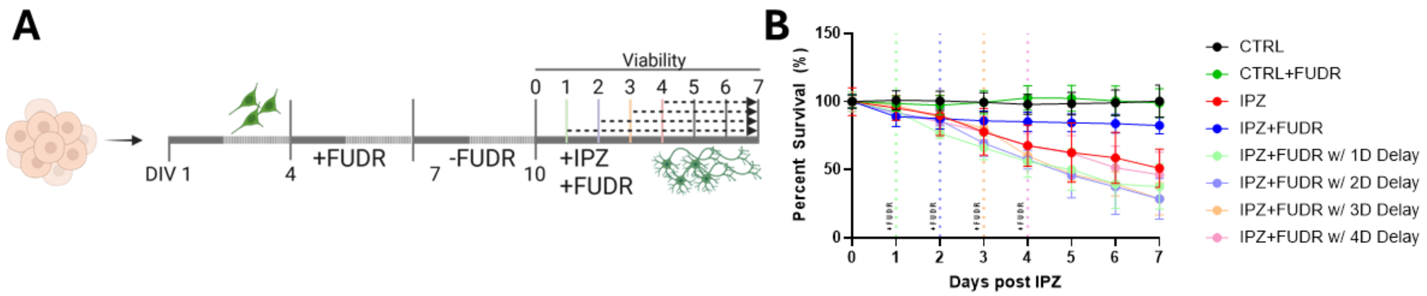


**Figure S7: Time-dependent viability curves with delayed FUDR treatments in iPSC-derived motor neurons post IPZ treatment. (A)** Timeline of iPSC-derived motor neuron generation treated with DMSO control or IPZ treatment with or without FUDR at DIV10, and analyzed by viability assays or imaging at the indicated time points post IPZ. **(B)** Seven-day viability curves of motor neurons treated with DMSO control or IPZ treatment with or without FUDR, analyzed by two-way ANOVA with Tukey's multiple-comparison test ($p < 0.05$) (n=3 replicates). Replicates are independent differentiations to account for batch variability. Schematics created with BioRender.com. All data points and statistical comparisons are provided in the Source Data file.

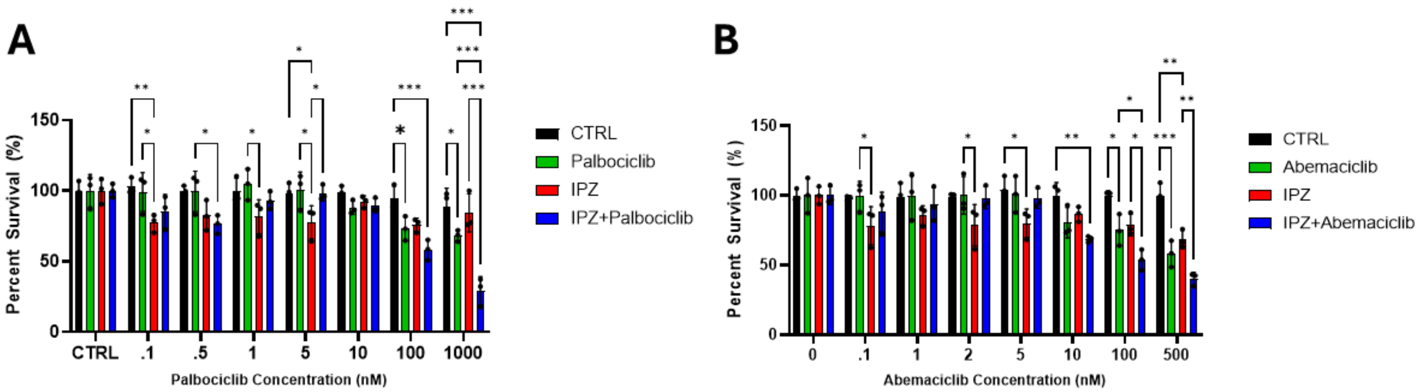


**Figure S8: Palbociclib and Abemaciclib dose-response curves (A-B)** Dose-response curve with and without importazole (10 µM) co-treated with increasing concentrations of palbociclib and abemaciclib from .1 nM to 1000 nM, analyzed by two-way ANOVA with Tukey's multiple-comparison test ($p < 0.05$) (n=3 replicates). Bar graphs represent mean ± standard deviation and significance values are indicated as * p<0.05, *** p<0.001. Replicates are independent differentiations to account for batch variability. All data points and statistical comparisons are provided in the Source Data file.
